# A multiscale spatiotemporal model including a switch from aerobic to anaerobic metabolism reproduces succession in the early infant gut microbiota

**DOI:** 10.1101/2021.08.22.457032

**Authors:** David M. Versluis, Ruud Schoemaker, Ellen Looijesteijn, Daniël Muysken, Prescilla V. Jeurink, Marcel Paques, Jan M. W. Geurts, Roeland M. H. Merks

## Abstract

The human intestinal microbiota starts to form immediately after birth, and is important for the health of the host. During the first days facultatively anaerobic bacterial species generally dominate, such as Enterobacteriaceae. These are succeeded by strictly anaerobic species, particularly *Bifidobacterium* species. An early transition to *Bifidobacterium* species is associated with health benefits: for example, *Bifidobacterium* species repress growth of pathogenic competitors and modulate the immune response. Succession to *Bifidobacterium* is thought to be due to consumption of intracolonic oxygen present in newborns by facultative anaerobes, including Enterobacteriaceae. To study if oxygen depletion suffices for the transition to *Bifidobacterium* species, here we introduce a multiscale mathematical model that considers metabolism, spatial bacterial population dynamics and cross-feeding. Using publicly available metabolic network data from the AGORA collection, the model simulates *ab initio* the competition of strictly and facultatively anaerobic species in a gut-like environment under the influence of lactose and oxygen. The model predicts that individual differences in intracolonic oxygen in newborn infants can explain the observed individual variation in succession to anaerobic species, in particular *Bifidobacterium* species. *Bifidobacterium* species become dominant in the model by using the bifid shunt, which allows *Bifidobacterium* to switch to sub-optimal yield metabolism with fast growth at high lactose concentrations as predicted here using flux-balance analysis. The computational model thus allows us to test the internal plausibility of hypotheses for bacterial colonization and succession in the infant colon.

**IMPORTANCE:** The composition of the infant microbiota has a great impact on infant health, but its controlling factors are still incompletely understood. The frequently dominant anaerobic *Bifidobacterium* species benefit health, e.g., because they can keep harmful competitors under control and modulate the intestinal immune response. Controlling factors could include nutritional composition and intestinal mucus composition, as well as environmental factors, such as antibiotics. We introduce a modeling framework of a metabolically realistic intestinal microbial ecology in which hypothetical scenarios can be tested and compared. We present simulations that suggest that greater levels of intra-intestinal oxygenation more strongly delay the dominance of *Bifidobacterium* species, explaining the observed variety of microbial composition and demonstrating the use of the model for hypothesis generation. The framework allows us to test a variety of controlling factors, including intestinal mixing and transit time. Future versions will also include detailed modeling of oligosaccharide and mucin metabolism.

## INTRODUCTION

The human infant microbiota starts forming directly after birth, and differs greatly from the adult microbiota (1). Three main community types are observed in the infant: (a) a Proteobacteria-dominated microbiota; (b) an Actinobacteria-dominated microbiota; and, less frequently (12-14% in one study (2)), (c) a Bacilli-dominated microbiota (1). The three main community types are established after an ecological succession of early communities. This ecological succession is thought to be controlled by nutrition and early oxygen in the colon. To develop hypotheses on the potential mechanisms and controlling factors of the initial development of the human microbiota, in particular the role of early oxygen, we introduce a computational model.

In the first 24 to 48 hours after birth, Proteobacteria, including *Escherichia coli* and *Enterobacter cloacae*, and Bacilli, including *Streptococcus, Lactobacillus*, and *Staphylococcus*, are the most common (3, 1). In the following weeks Proteobacteria are often replaced by anaerobic Actinobacteria, mainly *Bifidobacterium* species, whereas Bacilli are succeeded by either Proteobacteria or Actinobacteria. Other anaerobic species are also found, but typically do not dominate. Actinobacteria generally persist as the dominant group until weaning (1). A possible trigger for the replacement of the Enterobacteriacea by *Bifidobacterium* spp., is depletion of a hypothesized initial amount of oxygen (4, 5). Oxygen diffuses into the gut lumen from the body and is taken up by bacteria, by colonocytes, and by non-biological chemical processes in the cecal contents (6, 7, 8) leading to oxygen depletion. The relative importance of these processes is still under debate (7, 8).

*Bifidobacterium* species (1) are associated with a range of health benefits. For example, early succession to a *Bifidobacterium* dominated microbiota is correlated with reduced probabilities to be underweight at 18 months of age or to experience colic (2, 9). Acetate and lactate produced by *Bifidobacterium* also contribute to the acidification of the infant gut, which suppresses many pathogens (10). A key question, therefore, is if and how *Bifidobacterium* can be promoted. The composition and development of the infant microbiota is thought to be determined by many factors, including nutrition (11, 12, 13), i.e. human milk or infant formula. The presence of lactose, the most abundant carbohydrate in human milk and most infant formulas, is essential for acquiring high levels of *Bifidobacterium* (14). This suggests that nutrition can play a crucial role in promoting *Bifidobacterium*.

To obtain more insight into the potential ecological and metabolic mechanisms underlying bacterial colonization and succession in the infant colon, here we propose a multiscale, spatial computational model of the infant gut microbiota. Earlier work has introduced similar models of the adult gut microbiota (15, 16, 17, 18). These make use of flux balance analysis (FBA) to model metabolism. Given a number of constraints including substrate availability and enzyme availability, FBA uses genomescale metabolic network models (GEMs) to predict optimal biomass or energy production and the exchange of metabolites with the environment. In spatial FBA, FBA models representing small subpopulations of bacteria are coupled together to model a microbial, metabolizing ecosystem. While some approaches assume the whole system is in steady-state, allowing them to use the optimization approach for the whole ecosystem (16), here we build upon dynamic approaches such as BacArena (17), COMETS (19), and the approach by Van Hoek and Merks (18). In these models, the metabolite exchange rates given by FBA are used to dynamically change metabolite concentrations in the environment, and predict the resulting effects on the bacterial populations. These properties make these dynamic, spatial FBA approaches highly suitable for our purpose of modeling the formation of the infant gut microbiota.

To analyze succession of bacterial species in a dynamic ecosystem such as the infant gut, it is key to accurately describe the dependency of bacterial growth rate and metabolite exchange on environmental metabolite concentrations. Previous work has shown that these are well described by constraints on the metabolic capacity of bacterial cells. These are given by an ‘enzymatic constraint’ which poses a limit on the summed metabolic flux through the whole cell (20), or through more advanced methods such as flux balance analysis with molecular crowding (FBAwMC) (21) which weighs the flux constraint according to the efficiency and volume of the individual enzymes (22). Our model is, therefore, based upon the previous approach by Van Hoek and Merks (18) that uses FBAwMC. However, because such detailed data on enzyme efficiency and volume is unavailable for the majority of GEMs, here we have replaced FBAwMC with the simpler ‘enzymatic constraint’ approach (20). This approach allows us to model metabolic limitations and substrate-dependent switches in metabolism.

The model represents the first 21 days of development of the infant microbiota in a dynamic, spatially extended, gut-like environment, and considers three scales. At the microscopic scale, the model simulates individual metabolites. At the mesoscopic scale, metabolism and growth of bacteria occurs as a function of the local availability of metabolites. The local environment is depleted by the bacterial models, and receives the metabolites they deposit. This will influence the availability of metabolites in the next timesteps. The macroscopic scale represents the whole ecosystem. At this level a diffusion model simulates the spread of metabolites between adjacent lattice sites, and a local mixing model simulates mobility of bacterial populations. The metabolites also advect distally to represent luminal movements, while to represent adherence to the mucus the bacterial populations do not advect. Bacterial populations are also randomly deleted to represent local extinction, and they can form new populations in their immediate environment. The system currently considers 15 bacterial species based on 15 GEMs taken from the AGORA collection (23), a database of 773 semiautomatically reconstructed GEMs of human gut bacteria that includes all major infant species (23).

The infant gut microbiota is partially limited by carbohydrates. Prebiotic carbohydrates increase population sizes *in vitro* and in mice inoculated with infant gut bacteria (24, 25). The primary carbohydrate in human milk and nearly all infant formula is lactose, and infant formulas without lactose lead to a different infant microbiota with greatly reduced *Bifidobacterium* abundance (14). Therefore to a first approximation, we ignore key nutrient sources such as human milk oligosaccharides, fats, protein residues, or intestinal mucus and focus on carbohydrate metabolism, here taking lactose as the only input carbon source in the model.

In comparison with the previous approach (18) upon which the present work builds, the technological advance is that the present framework simulates a dynamic ecology of a large diversity of bacterial species. To this end, the modeling framework can, with minor modifications, import any GEM written in SBML, such as those in the AGORA database that we use in this project (23). The previous system (18) could only describe a ‘superbacterial’ metabolic network representative of the adult microbiota, while bacterial diversity was simulated by activating or repressing individual pathways. In the present work we study a selection of species representative of the infant microbiota. The new system also allows us to analyze and visualise the flow of fluxes through the emergent network of dynamically interacting spatially distributed populations. The conceptual advance following from this technological advance is that we can now predict the effect of environmental conditions (oxygen, food, etc.) on the relative abundance of individual bacterial species. This advance has allowed us to use the system to demonstrate the mechanistic consistency of the hypothesis (4, 5) that a transition from an Enterobacteriaceae-dominated microbiota to a *Bifidobacterium* spp.-dominated microbiota is explained through gradual depletion of oxygen leading to anaerobiosis. The model suggests that high levels of initial oxygen can lead to prolonged dominance by Enterobacteriaceae, even after oxygen has been depleted, possibly explaining observed variation in the composition of the infant microbiota (1, 13), and reveals possible cross-feeding interactions between the species. Altogether the model shows to be a useful tool for qualitatively evaluating hypotheses on the dynamics of the infant gut microbiota.

## RESULTS

### Model outline

The model of the infant colon is discussed in detail in Section Methods. Briefly, given a typical infant colon length of 45 cm (26) and diameter of approximately 1.5 cm (27) the colon is described on a regular square lattice of 225 × 8 boxes of 2 *mm* × 2 *mm* each, thus reaching a good balance between computational efficiency and accuracy (Fig. 1A). Because bacterial populations can vary at a much smaller scale (28, 29), the boxes are purely used as a computational discretization. Each lattice site can contain a population of one species, representing only the locally dominant bacterial species. It is described by a GEM from a list of 15 species prevalent in the earliest infant gut microbiota ((30, 11) selected from the AGORA collection (23)) (Table 1, Methods section Species composition).

**TABLE 1.**
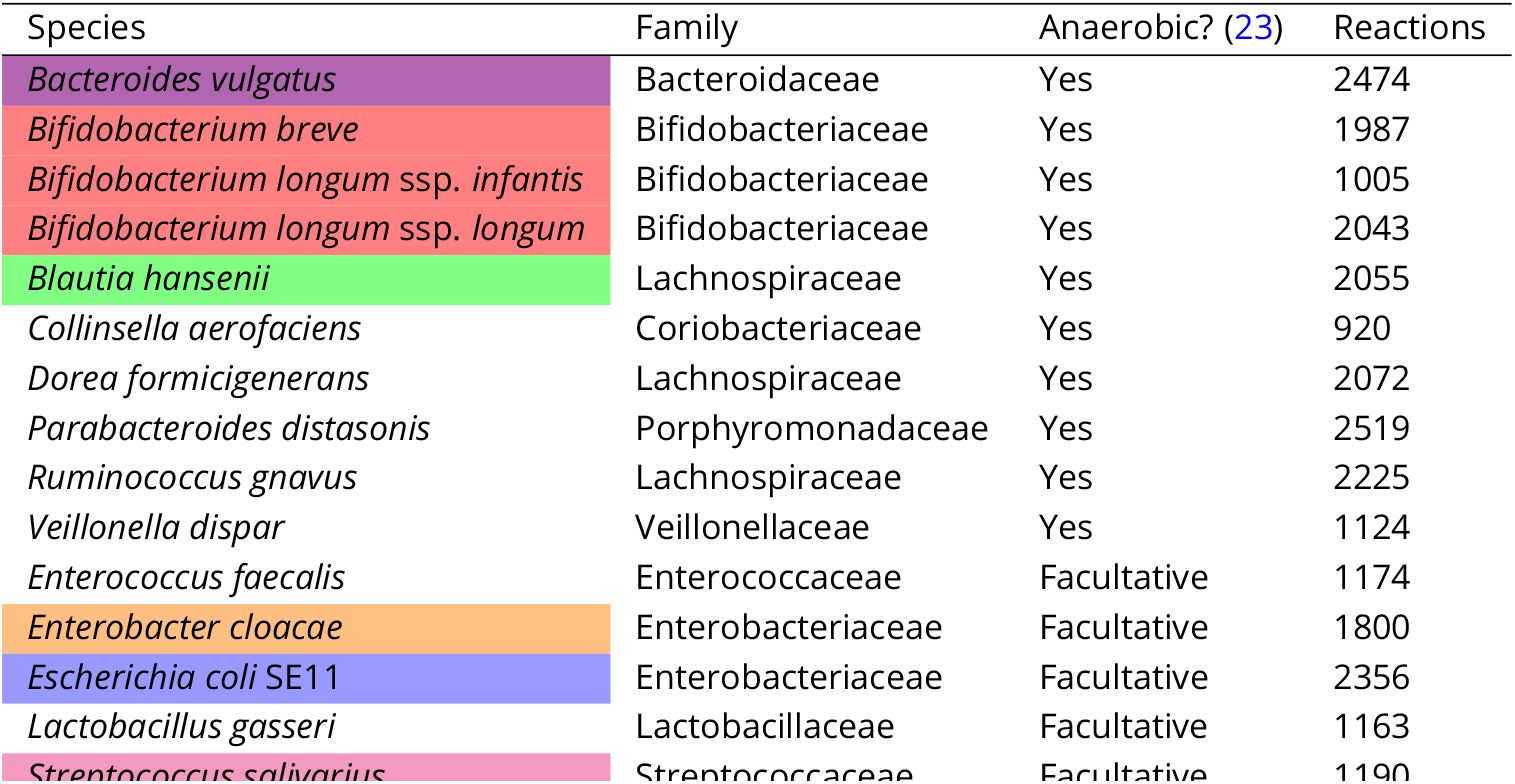
Species included in our model, all from the AGORA collection(23). Color indicates color used in figures

| Species | Family | Anaerobic? (23) | Reactions |
| --- | --- | --- | --- |
| *Bacteroides vulgatus* | Bacteroidaceae | Yes | 2474 |
| *Bifidobacterium breve* | Bifidobacteriaceae | Yes | 1987 |
| *Bifidobacterium longum* ssp. *infantis* | Bifidobacteriaceae | Yes | 1005 |
| *Bifidobacterium longum* ssp. *longum* | Bifidobacteriaceae | Yes | 2043 |
| *Blautia hansenii* | Lachnospiraceae | Yes | 2055 |
| *Collinsella aerofaciens* | Coriobacteriaceae | Yes | 920 |
| *Dorea formicigenerans* | Lachnospiraceae | Yes | 2072 |
| *Parabacteroides distasonis* | Porphyromonadaceae | Yes | 2519 |
| *Ruminococcus gnavus* | Lachnospiraceae | Yes | 2225 |
| *Veillonella dispar* | Veillonellaceae | Yes | 1124 |
| *Enterococcus faecalis* | Enterococcaceae | Facultative | 1174 |
| *Enterobacter cloacae* | Enterobacteriaceae | Facultative | 1800 |
| *Escherichia coli* SE11 | Enterobacteriaceae | Facultative | 2356 |
| *Lactobacillus gasseri* | Lactobacillaceae | Facultative | 1163 |
| *Streptococcus salivarius* | Streptococcaceae | Facultative | 1190 |

**FIG 1.**
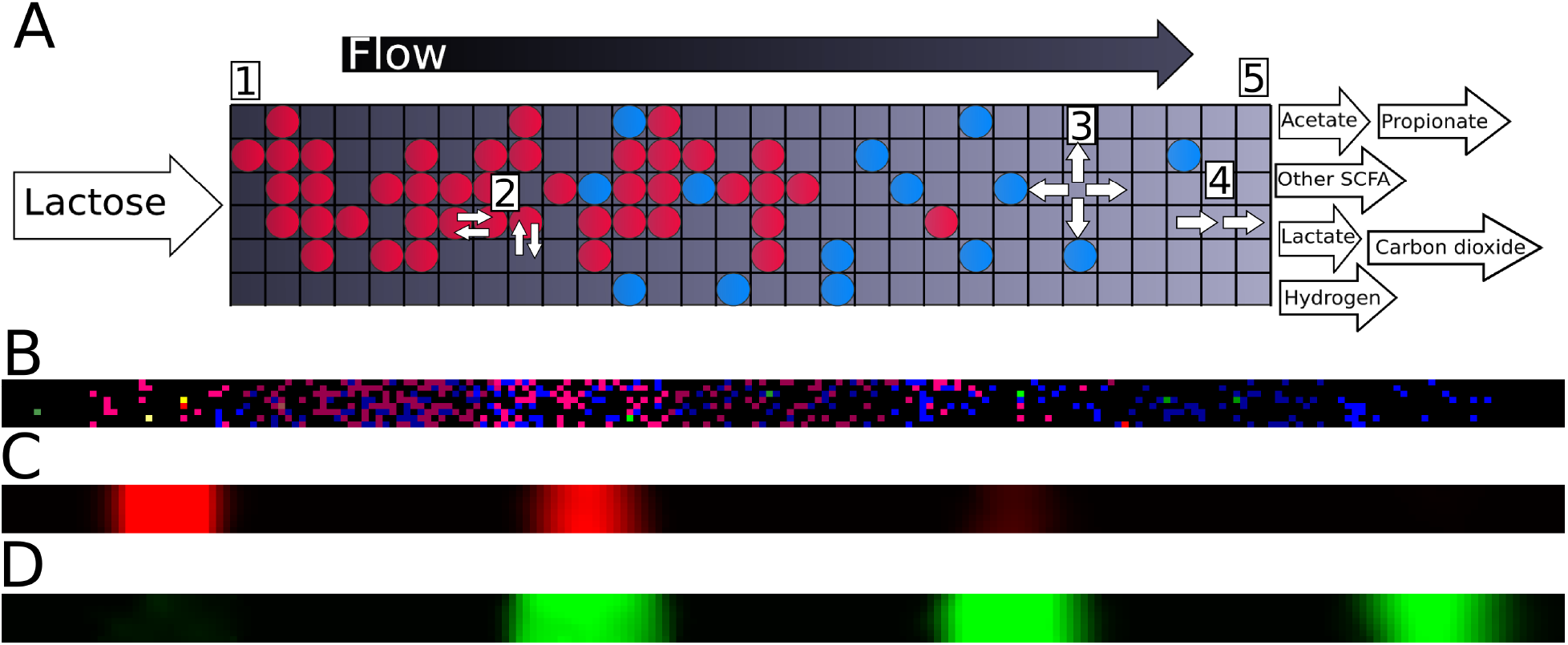
The multi-scale metabolic model (A) Schematic of the model at a system level. Circles represent bacterial populations, color represents species. Flow through the tube is from left (proximal) to right (distal). Lactose is placed at the proximal side. Output metabolites are examples, and depend on bacterial metabolism. Lattice dimensions and ratio are schematic. (B) Screenshot of the bacterial layer of the model at a single time point. Color indicates species, brightness indicates growth rate on the current timestep. (C,D) Screenshots showing, respectively, lactose and lactate concentration of the model corresponding to B. Intensity indicates concentration.

A simulation proceeds as follows. We initialize the system by placing approximately 540 small populations of the 15 species randomly across the lattice (Fig. 1B). We run the model in timesteps representing three minutes of real time, these timesteps proceed as follows: To mimic carbohydrates entering the colon from the small intestine, a pulse of lactose is introduced every 60 timesteps to the lattice sites of the six most proximal columns (Fig. 1A-1). Then we predict the metabolism of local bacterial populations using FBA. To mimic metabolic trade-offs, we use an enzymatic constraint that sets the maximum summed flux for each FBA solution. The uptake bounds for each population are set according to the local concentrations of metabolites (Fig. 1C&D). FBA returns a solution that maximizes the rate of ATP production, which is a suitable proxy for the rate of biomass production (31) given the focus on carbohydrate metabolism. This biomass production rate (indicated by brightness in Fig. 1B) is used to update the local population size. Populations spread into an available adjacent lattice site once the population size has exceeded a threshold. The FBA solution also returns a set of metabolite exchange rates, which are used to update the local metabolite concentrations (Fig. 1D). In total, 723 extracellular metabolites are described in the model, but only 17 of these are typically found in more than micromolar amounts in our simulations (Methods, Table 2). Bacterial populations diffuse through swaps of the content of adjacent lattice sites (Fig. 1A-2). Metabolites diffuse between adjacent lattice sites (Fig. 1A-3) and advect to more distal sites (Fig. 1A-4) over one lattice per timestep, reaching transit times of effectively 11 hours (32, 33). Metabolites and populations that reach the distal side of the system (Fig. 1A-5) are removed. To mimic the appearance of new bacterial populations (e.g. from ingested spores or introduction from intestinal crypts (34)) each empty lattice site now has a small probability to be filled with a new bacterial population selected at random from all available species (Table 1). This completes the timestep, after which we record the metabolite concentrations and bacterial population in the whole model. All parameters are given in Section Parameters. We have also performed a sensitivity analysis for several parameters around the default parameter set shown in Table 2 (Section Sensitivity analysis).

**TABLE 2.** Extracellular metabolites present in the model in more than micromolar amounts

| Common name | BiGG ID |
| --- | --- |
| Acetate | ac |
| Alpha-Ketoglutaric acid | akg |
| Carbon dioxide | co2 |
| Ethanol | etoh |
| Formate | for |
| Hexadecanoate | hdca |
| Hydrogen | h2 |
| L-Lactate | lac_L |
| D-Lactate | lac_D |
| Lactose | lcts |
| Octadecanoate | ocdca |
| Oxygen | o2 |
| Propionate | ppa |
| Proton | h |
| Succinate | succ |
| Tetradecanoate | ttdca |
| Water | h2o |

We performed several checks to ensure the biochemical and thermodynamic plausibility of the model (Methods section Checking the validity of the GEMs). After the resulting corrections to the GEMs (S1 Table), no atoms were generated or lost by the FBA solutions, there was no flux through the objective function without a substrate, and 99.999% of all fluxes of Fig. 3A contained less free energy in the output than in the input. The remaining 0.001% all had no energy increase and growth rates less than 0.001% of the average growth rate. We therefore concluded that the system conserved mass and energy.

**FIG 2.**
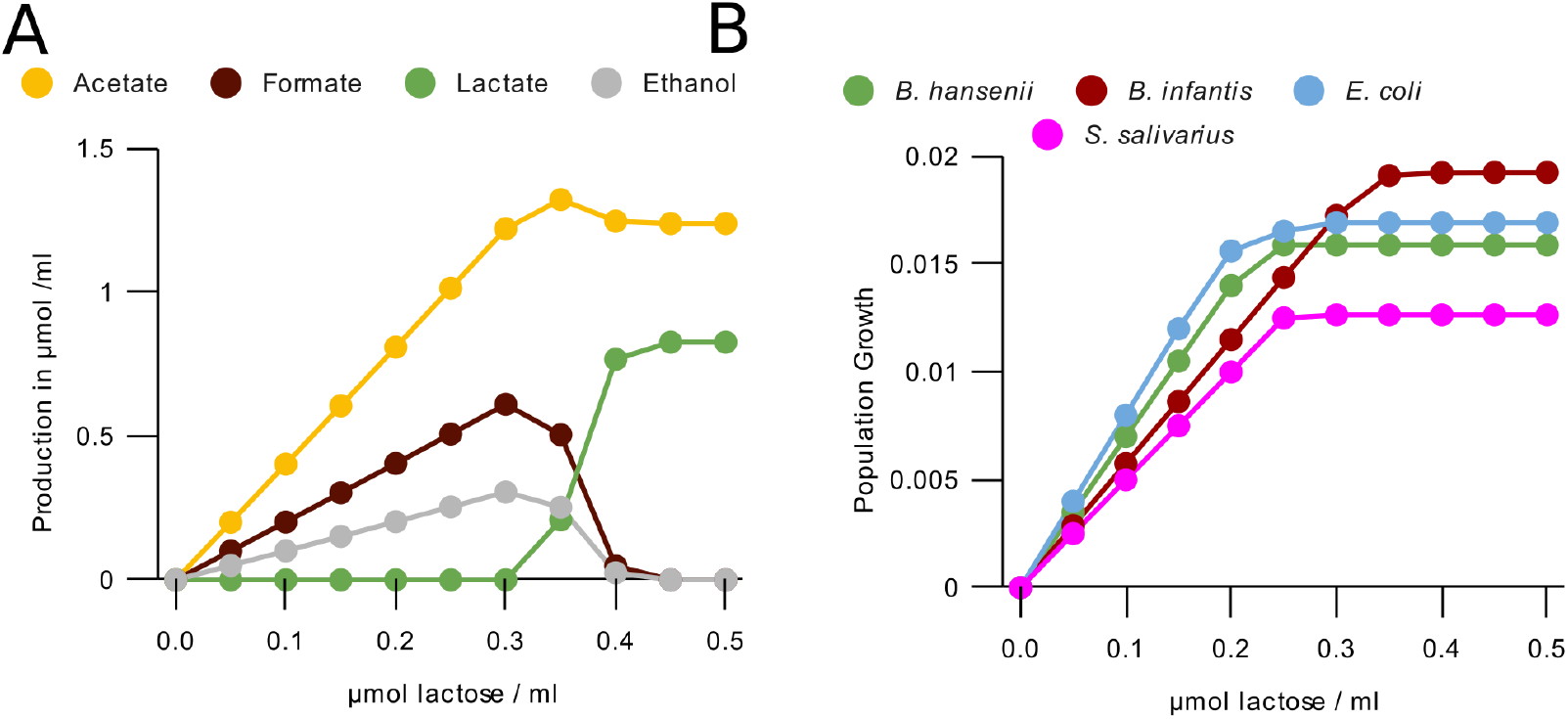
Enzymatic constraint reproduces metabolic switching in *Bifidobacterium* (A) Production of metabolites per timestep by a *B. longum infantis* population of 5 · 10^9^ bacteria with access to one lattice site (0.05ml) using our FBA with enzymatic constraint method. (B) Growth per timestep by lactose concentration for populations of 5 · 10^9^ bacteria with access to one lattice site (0.05ml) of some major bacterial species using our FBA with enzymatic constraint method. *B. longum infantis* is used as the *Bifidobacterium* representative.

**FIG 3.**
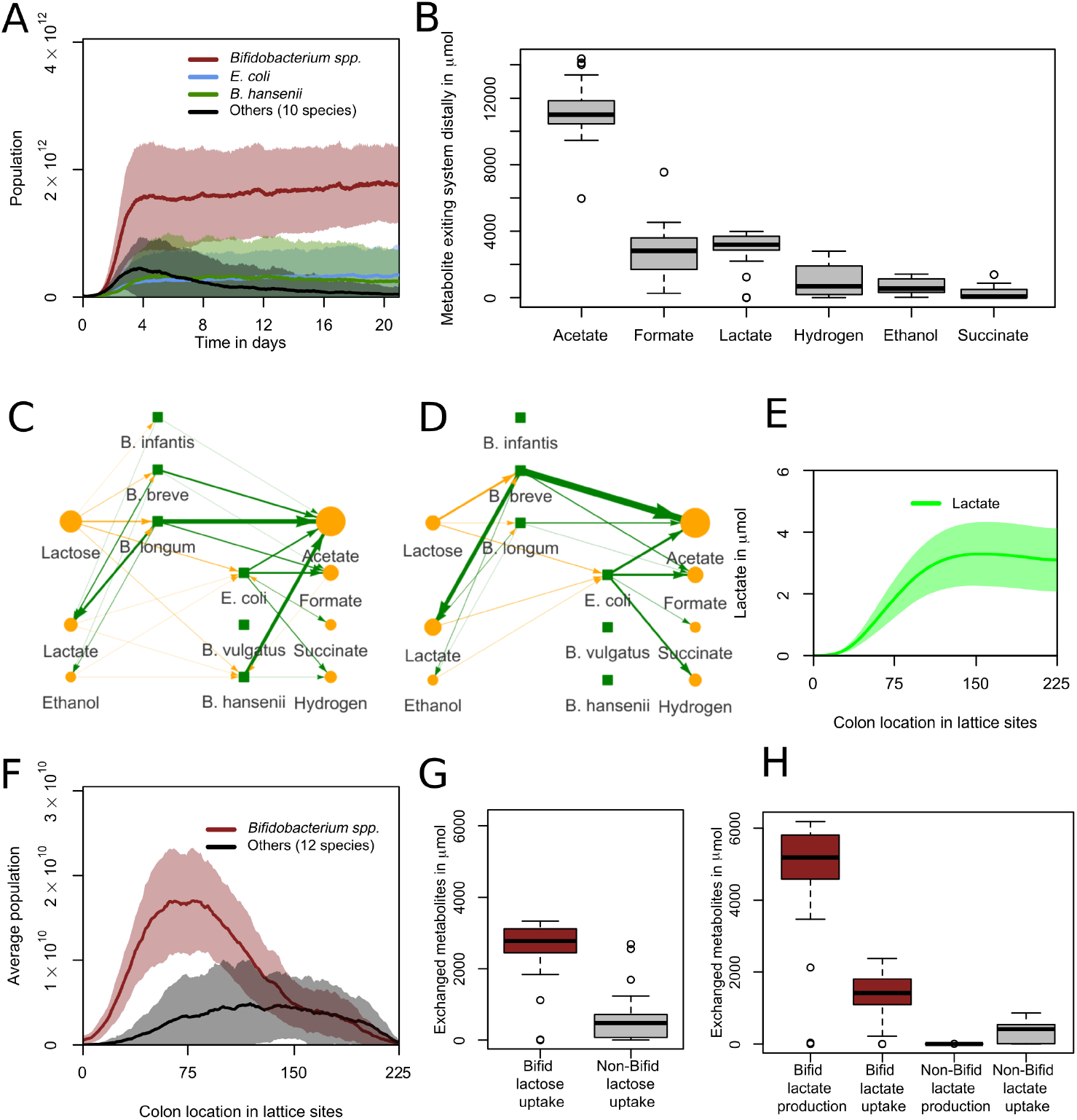
*Bifidobacterium* spp become dominant anaerobically and mostly proximally with a unique metabolic profile (A) Abundances for grouped species over 21 days. One day is 480 timesteps. Curve shows mean value and shaded area one standard deviation over n=30 simulations. (B) Distribution of metabolites exiting the system distally over the last two days (960 timesteps) of the simulation. n=30. (C,D) Visualisation of metabolic interactions in the sample run of S1 Video. Green lines represent production, yellow lines represent consumption. Line width and intensity is proportional to the amount exchanged with the environment, with a threshold of 0.5 µmol, with no normalization. Metabolite circle size is relative to the most abundant metabolite, with a minimum displayed size of 26 pixels. Data from hour 90 to 93 (step 1800 to 1860) for C and last 3 hours (step 10020 to 10080) for D (E) Spatial distribution of lactate over the last two days (960 timesteps) of the simulation. Curve shows mean value and shaded area one standard deviation over n=30 simulations. (F) Abundance per location for grouped species over the last two days (960 timesteps). Curve shows mean value and shaded area one standard deviation over n=30 simulations. (G) Metabolism of lactose by grouped species. All data from last two days. n=30. (H) Metabolism of lactate by grouped species. All data from last two days. n=30.

### Enzymatic constraint reproduces metabolic switching in *Bifidobacterium*

Many microorganisms switch between metabolic pathways of higher and lower yield depending on substrate availability. These so called metabolic switches can be reproduced with FBA through an enzymatic constraint (20, 35, 36, 22) that limits the summed metabolic flux of a bacterium. *Bifidobacterium* also displays a metabolic switch through a pathway known as the bifid shunt. At low extracellular sugar concentrations, *Bifi-dobacterium* produces a mixture of acetate, ethanol, and formate in a 1:1:2 ratio from pyruvate. At increased sugar concentrations the bifid shunt instead reroutes pyruvate to lactate. In addition acetate is produced from acetyl-phosphate independently, such that at low concentration of sugars *Bifidobacterium* produces acetate, ethanol and formate in a 4:1:2 ratio, and at increased sugar concentrations *Bifidobacterium* switches to production of acetate and lactate in a 3:2 ratio (37, 38).

To test if the enzymatic constraint suffices to explain the metabolic switch in *Bifidobacterium*, we performed simulations of a single timestep in increasing lactose concentrations (Fig. 2A). These correctly predicted the metabolic switch. The ratios between the metabolites before and after the metabolic switch, as well as the transient between the two regimes and the pathways used (S2 Table) match the experimental observations. Simulations without the constraint only reproduced the situation at low concentrations of lactose (S1 Figure A), showing that the enzymatic constraint is required. The model also predicts an unobserved metabolic switch on lactose in *E. coli* (S1 Figure B) and reproduced the metabolic switch in *Bacteroides vulgatus* (39) (S1 Figure C). Another effect of the enzymatic constraint is that the growth rate saturates at a species-dependent concentration (Fig. 2B, S1 Figure D). As a consequence the relative growth rates of the species depend on the lactose concentration.

### Model predicts *Bifidobacterium* dominance through metabolism under anaerobic conditions

After having established the effect of the enzymatic constraint in models of individual bacterial populations, we studied the behavior of the full multiscale model for anaerobic conditions. After 21 days (10080 timesteps) *Bifidobacterium* spp. had become the most abundant in 27 of 30 simulations (Fig. 3A, S1 Video), a typical situation found in infants of a corresponding age (11, 2). Also in agreement with *in vivo* data, the model predicted the presence of populations of *E. coli*. Our model also predicted populations of *Blautia hansenii*, which is only occasionally present in the infant colon (11). We will investigate the niche of this species further in the section ‘Non-*Bifidobacterium* species benefit from lactate consumption’. All species other than the aforementioned had abundances either lower or not significantly higher than their initial abundance after 21 simulated days (p>0.05, two-sample t-test). To test if these species were required for the formation of the community we ran an additional 30 simulations in which we only included species that had an abundance significantly larger than their initial abundance at the end of other simulations. This led to a very similar set of outcomes (S2 Figure A). In these simulations none of the species abundances differed from the original simulations with all species included (p>0.05, two-sample t-test). In the remaining simulations all species are included in order to prevent an initial bias, thus leaving in the possibility that any of the rare species become abundant under one of the conditions considered. In practice this did not occur.

We next analysed the metabolites diffusing or advecting out of the distal end of the gut over the last two days of the simulations of Fig. 3A. We found considerable amounts of acetate, formate, lactate, hydrogen, ethanol, and succinate (Fig. 3B). All other metabolites were present in much smaller quantities. The high abundance of acetate and lower abundance of lactate agree with the fecal composition of three month old formula-fed infants (40), and is characteristic of microbiota dominated by *Bifidobacterium*. Formate, hydrogen, ethanol and succinate are also all reported in infant feces (41, 42, 43, 44).

To examine how this metabolic pattern is formed we analysed the metabolic production and uptake in a sample run (Fig. 3C). *Bifidobacterium* converts lactose into a mix of acetate, lactate, ethanol, and formate, as predicted (Fig. 2A), with a higher lactate production than ethanol or formate. Some of this lactate is reabsorbed in later timesteps and converted into the three other metabolites by *Bifidobacterium*. A mixture of lactose and crossfeeding products is converted by *E. coli* and *B. hansenii* into the remaining metabolites of Fig. 3B. The main crossfeeding nutrient absorbed is lactate, produced by *Bifidobacterium*, but there is also uptake of ethanol, formate, acetate, and succinate. These are also produced by the crossfeeders. The network has become simpler at the end of this run, as *B. hansenii* is at a very low abundance and *E. coli* now only consumes lactose, lactate, and ethanol, which it does not produce (Fig. 3D).

### Crossfeeding emerges in simulations

The observed crossfeeding in the sample run (Fig. 3C&D) suggested that production of metabolites by *Bifidobacterium* spp. may be crucial for *E. coli* and *B. hansenii*. To further analyse such crossfeeding interactions we next analysed the spatial distribution of lactose and the Bifidobacterial products lactate and acetate. From day 19 to 21, lactose was present mostly proximally (S2 Figure B), lactate was most abundant in the middle of the colon (Fig. 3E), and acetate was most abundant distally (S2 Figure C). To analyse the role of *Bifidobacterium* spp. in shaping this metabolic pattern, we analysed the location of populations from day 19 to 21 (Fig. 3F). *Bifidobacterium* spp. was located proximally, whereas the other species were located more distally. These results suggested that separated metabolic niches had formed in the simulated colon. To quantify crossfeeding interactions we summed the flux of lactose and lactate for *Bifidobacterium* and for all non-*Bifidobacterium* species grouped together. *Bifidobacterium* spp. was the largest consumer of lactose (Fig. 3G) and both the largest producer and consumer of lactate (Fig. 3H). An additional consumption of lactate towards acetate, formate and ethanol takes place at lower lactose concentrations, when the enzymatic constraint is not saturated by lactose uptake. However, the non-*Bifidobacterium* groups consumed more lactate relative to their lactose consumption. To test the effect of lactate uptake by *Bifidobacterium*, additional simulations were run in which the conversion from lactate to pyruvate was blocked, preventing *Bifidobacterium* from consuming lactate. In these simulations *Bifidobacterium* was the most abundant group in 20 of the 30 simulations, with the largest average abundance (S2 Figure D). Thus lactate consumption played a small role but was not the primary cause of *Bifidobacterium*’s dominance. We hypothesized that *Bifidobacterium*’s primary niche is as primary consumer, consuming lactose, while for other species niches exist as primary and/or secondary consumer around consuming lactose or lactate. Combined with the observation (Fig. 3E) that lactate concentration is lower in the distal colon and the recorded crossfeeding fluxes in our network visualisation (S1 Video), we can conclude that the population at the distal end of the simulated colon consists at least partially of such secondary consumers, i.e. crossfeeders.

### Lactose uptake through the bifid shunt is essential for *Bifidobacterium* dominance

Next we determined whether the bifid shunt is essential to *Bifidobacterium* dominance in the model. We disabled the bifid shunt by blocking all flux through the reactions associated with fructose-6-phosphate phosphoketolase and ran 30 simulations. Fructose-6-phosphate phosphoketolase is unique to and necessary for the bifid shunt, and characteristic of *Bifidobacterium* as a genus (45). *Bifidobacterium* spp. were dominant in none of these simulations (Fig. 4A, S2 Video). Instead, *E. coli, B. vulgatus*, and *B. hansenii* were dominant. This indicates that the bifid shunt is crucial to the dominance of *Bifidobacterium* spp. in our model. We also ran 30 simulations in which only the production of lactate was disabled in *Bifidobacterium* (S2 Figure F). 27 of these simulations still led to *Bifidobacterium* dominance. This indicates that the metabolic switch to lactate production was not essential to *Bifidobacterium* dominance in our model. *B. vulgatus* also had a higher average abundance, but this was not significant (p=0.27, two-sided t-test).

**FIG 4.**
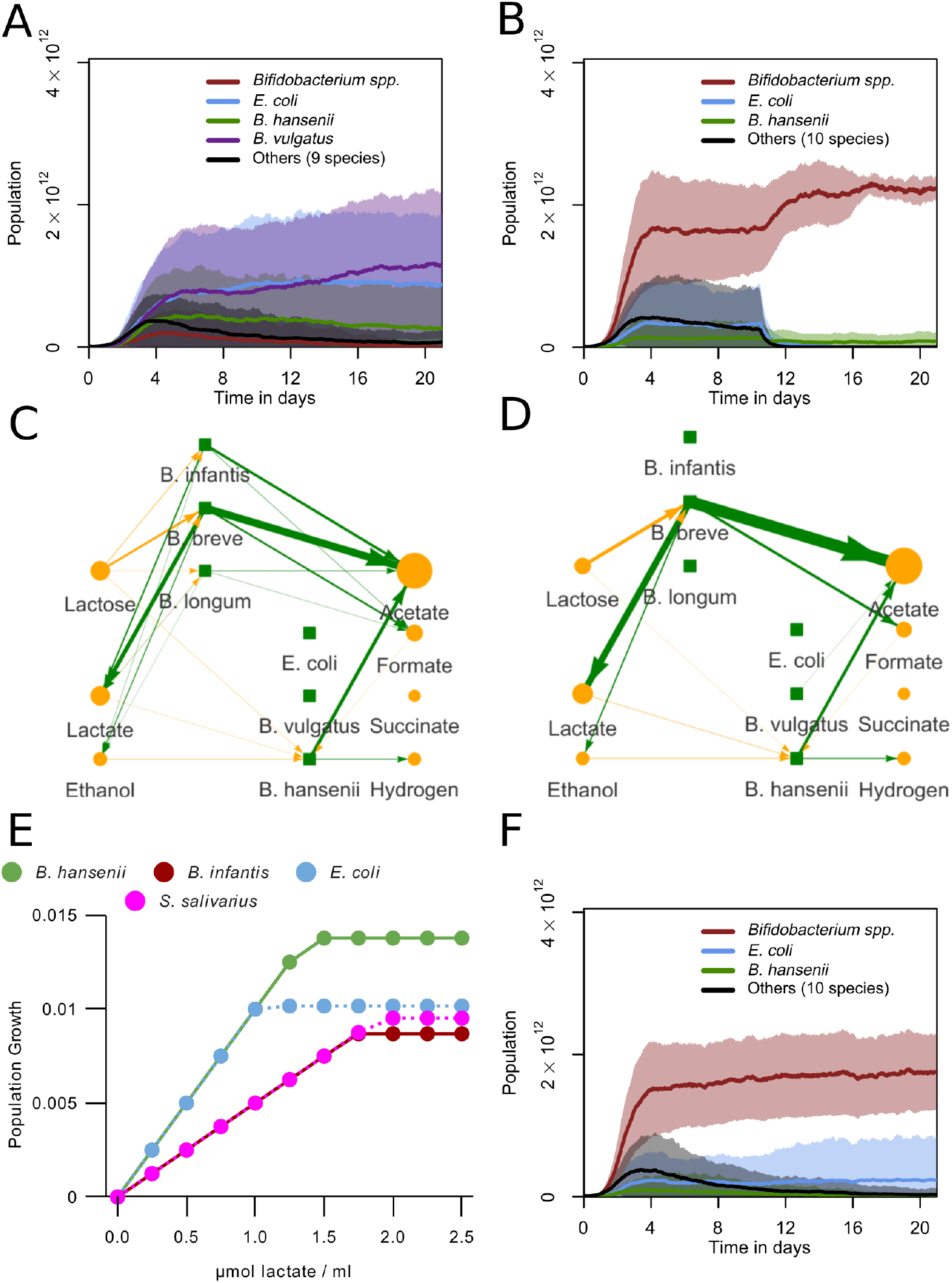
*Bifidobacterium* and *E. coli* require lactose, but *B. hansenii* requires lactate (A) Abundances for grouped species over 21 days. The bifid shunt was disabled for all *Bifidobacterium* species. One day is 480 timesteps. Curve shows mean value and shaded area one standard deviation over n=30 simulations. (B) Abundances for grouped species over 21 days, but with lactose disabled for non-*Bifidobacterium* species from 5040 timesteps (10.5 days) onwards. One day is 480 timesteps. Curve shows mean value and shaded area one standard deviation over n=30 simulations. (C,D) Visualisation of metabolic interactions in a sample run. Green lines represent production, yellow lines represent consumption. Line width and intensity is proportional to the amount exchanged with the environment, with a threshold of 0.5 µmol, with no normalization. Metabolite circle size is relative to the most abundant metabolite, with a minimum displayed size of 26 pixels. Data from hour 90 to 93 (step 1800 to 1860) for C and last 3 hours (step 10020 to 10080) for D. (E) Growth per timestep by lactate concentration for populations 5 · 10^9^ bacteria with access to one lattice site (0.05ml) of some major bacterial species (F) Abundances for grouped species, with lactate consumption disabled for all species. One day is 480 timesteps. Curve shows mean value and shaded area one standard deviation over n=30 simulations.

### *B. hansenii* but not *E. coli* can sustain itself on crossfeeding

We observed that non-*Bifidobacterium* species appeared in the simulation behind a proximal population of *Bifidobacterium* spp. (Fig. 3F). These non-*Bifidobacterium* species consumed lactose (Fig. 3D&G), despite the fact that lactose concentrations have dropped compared to the proximal part (S2 Figure B). We therefore wondered how these non-*Bifidobacterium* species could persist in the model. At reduced lactose concentrations, *E. coli* produces more ATP per timestep than *Bifidobacterium* spp. (Fig. 2B). We hypothesized that non-*Bifidobacterium* species outcompete *Bifidobacterium* at reduced lactose concentrations in the model. To test this hypothesis, we blocked lactose consumption by non-*Bifidobacterium* species after 10.5 days, the midway point of the simulation, and ran 30 simulations. In this way, we ensured that sufficient secondary resources were produced by *Bifidobacterium* in case non-*Bifidobacterium* species made use of them. We observed a sharp population decrease of non-*Bifidobacterium* species starting directly after blocking lactose uptake (Fig. 4B). *E. coli* had a near-zero abundance at the end of these simulations. *B. hansenii* also had a significantly lower abundance compared to the baseline (p<0.01, two-sample t-test), but still had a presence (S2 Video). Thus, in our simulations, some amount of primary consumption was essential for *E. coli*, but not for *B. hansenii*.

### Non-*Bifidobacterium* species benefit from lactate consumption

To analyse how *B. hansenii* could sustain a population as secondary consumer, we investigated the substrates used by non-*Bifidobacterium* species in the model. Analysis of the flow of metabolites between species in a new run showed that the uptake of lactose by *B. hansenii* decreases over time, while the uptake of lactate increased (Fig. 4C&D). *E. coli* and *B. hansenii* also produce more ATP from lactate than *Bifidobacterium* at any concentration (Fig. 4E). Combined with our earlier observations of crossfeeding on lactate (Fig. 3D&H) we hypothesized that *Bifidobacterium* species cannot compete with *E. coli* and *B. hansenii* on pure lactate. Consumption of lactate by *B. hansenii* could be essential to its ability to sustain populations without lactose uptake.

To investigate if lactate is essential in feeding secondary consumer populations, we blocked the lactate uptake reaction for all species and ran 30 simulations. (Fig. 4F, S2 Video). At 21 days *B. hansenii* populations were much smaller than in simulations with functional lactate uptake, not significantly larger than the initial populations (p>0.05, two-sample t-test). *E. coli* populations were also significantly smaller than those in simulations with lactate uptake, but also significantly larger than their initial population at the start of the run (p<0.05, two-sample t-test). *Bifidobacterium* populations were similar to those in the baseline simulations (Fig. 3A). Together with the results in Fig. 4B this indicates that *E. coli* does not require lactate as a crossfeeding substrate like *B. hansenii* does.

### *E. coli* outcompetes *Bifidobacterium* by taking up early oxygen

During the first days Enterobacteriaceae or Bacilli such as *Streptococcus* are dominant in the *in vivo* infant gut microbiota, after which *Bifidobacterium* spp. are often dominant (1, 11). This pattern may be explained by the presence of oxygen in the gut shortly after birth (5, 3), e.g. via diffusion through the gut wall (7) or accumulation in the newborn infant gut *in utero* in the absence of microbes (46). Intracolonic oxygen diffusion depends on host tissue oxygenation and colonic blood flow (47, 48). To mimic the early presence of oxygen we introduced 0.1 to 10 µmol initial oxygen per lattice site. These values were chosen arbitrarily in absence of precise data, but cover a wide range of outcomes in our model. The oxygen diffused, but did not advect in the model.

0.1 µmol initial oxygen sufficed to reproduce the pattern observed *in vivo*: *E. coli* initially dominated and *Bifidobacterium* became dominant after a few days or weeks of simulated time (Fig. 5A, S3 Video). However, we did not see dominance of *Streptococcus* in any of the simulations, in contrast to observations *in vivo* (2). The initial presence of oxygen strongly affected the other species in the model: *B. hansenii* remained nearly entirely absent, while *B. vulgatus* had a higher abundance than it did without initial oxygen. In the network analysis of a sample run a similar pattern occurs as in the networks of Fig. 3C, but with different early species (Fig. 5B). Here the Enterobacteriaceae *E. coli* and *Enterobacter cloacea* consume more of the early lactose, along with oxygen. By the end of the run *Bifidobacterium* spp. again took up the role of primary lactose consumer and lactate producer, whereas *E. coli* had fed largely on lactate (Fig. 5C), thus assuming a role of secondary consumer.

**FIG 5.**
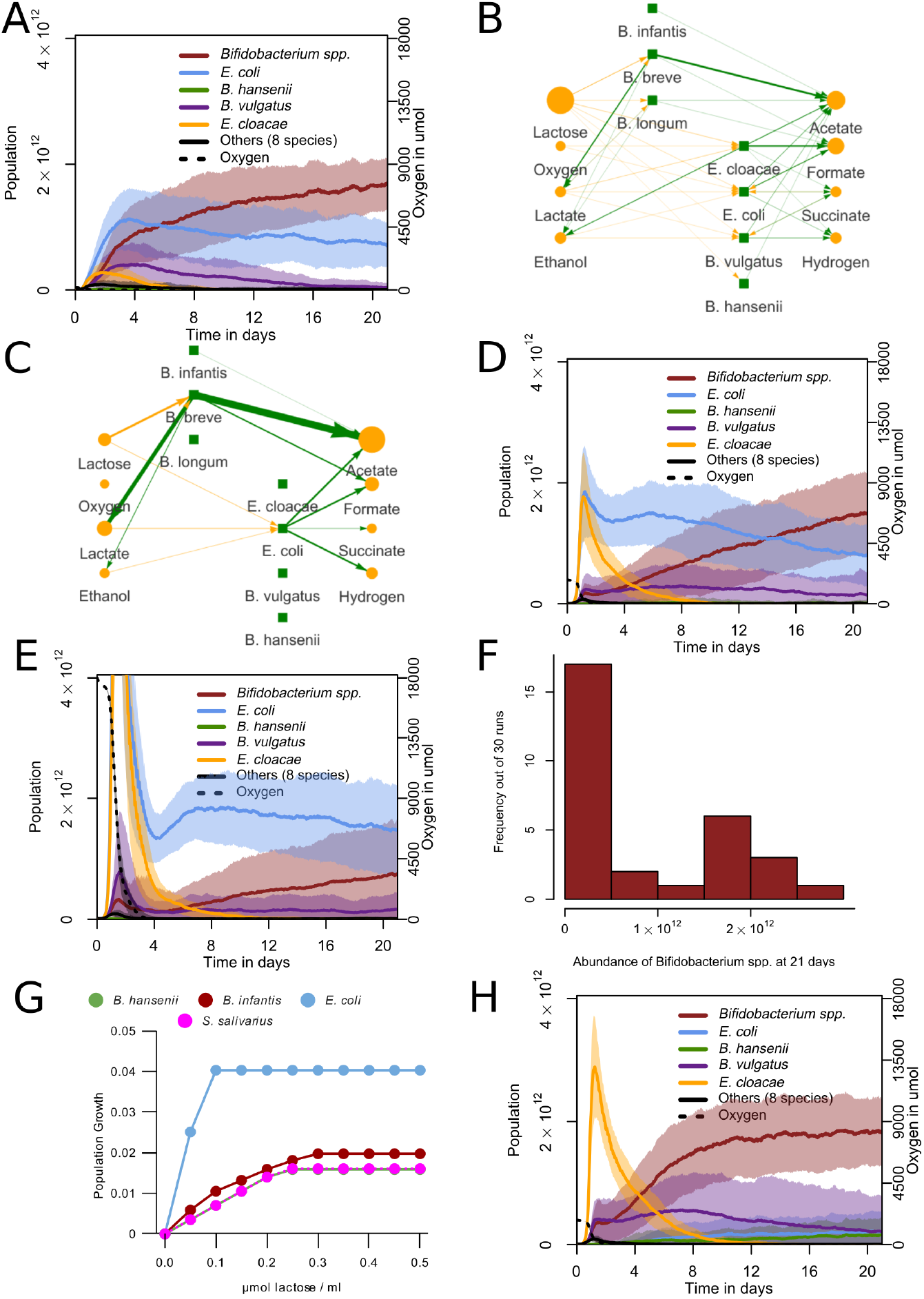
Initial oxygen leads to initial *E. coli* dominance and succession by *Bifidobacterium* (A,D,E) Abundances for grouped species. One day is 480 timesteps. Curve shows mean value and shaded area one standard deviation over n=30 simulations. Initial oxygen per lattice site in µmol is (A) 0.1 (D) 1 (E) 10 (B,C) Visualisation of metabolic interactions in a sample run. Green lines represent production, yellow lines represent consumption. Line width and intensity is proportional to the amount exchanged with the environment, with a threshold of 0.5 µmol, with no normalization. Metabolite circle size is relative to the most abundant metabolite, with a minimum displayed size of 26 pixels. Data from hour 30 to 33 (step 600 to 660) for B and last 3 hours (step 10020 to 10080) for C (F) Distribution of total *Bifidobacterium* abundance at 21 days (10080 timesteps) performed with 10 µmol initial oxygen per lattice site n=30. (G) Growth per timestep by lactose concentration in the presence of abundant oxygen for populations of 5 · 10^9^ bacteria with access to one lattice site (0.05ml) of some major bacterial species (H) Abundances for grouped species with oxygen uptake disabled for *E. coli*. Curve shows mean value and shaded area one standard deviation over n=30 simulations.

At 1 or 10 µmol initial oxygen per lattice site *E. coli* and *E. cloacea* remained dominant and the main primary consumers for longer (Fig. 5D, S4 Video), and succession to *Bifidobacterium* took much longer, or did not occur at all within the simulated timeframe (Fig. 5E). At 10 µmol oxygen per lattice site *Bifidobacterium* either dominated the population or remained much smaller, with few intermediate cases, leading to a bimodal distribution (Fig. 5F). This prediction matches *in vivo* observations (1, 13). The proportion of infants dominated by Enterobacteriaceae at 21 days of age varies depending on the study population, but can be 22% (2) or 25% (49) after 21 days, an intermediate value between the 10% (Fig. 5D) and the 60% (Fig. 5E) that we observe with different oxygen levels. We also initialised four sets of simulations in which oxygen was released from the upper and lower boundaries of the system every timestep (S3 Figure A-D). This led to an increasingly stronger stimulation of *E. coli* with higher levels of oxygen, but did not lead to any of the temporal effects observed *in vivo*. In a separate set of simulations we stopped oxygen release at the midpoint of our simulations. We observed an increase of *Bifidobacterium* spp. and a decrease of *E. coli* after that point (S3 Figure E), leading to a bimodal outcome similar to the simulations with initial oxygen (S3 Figure F).

We further examined the causes of *E. coli* dominance over *Bifidobacterium* in the presence of oxygen in our model. Fig. 5G shows that when oxygen is available *E. coli* has a much higher growth per timestep in our model than *Bifidobacterium* spp. for all concentrations of lactose, even though *Bifidobacterium* spp. also produce more ATP than under anaerobic conditions. *B. hansenii* and *Streptococcus salivarius*, in contrast, grew slower than *Bifidobacterium* for all concentrations in the presence of oxygen. This effect depends on the local oxygen concentration, but *E. coli* grows faster than other species even at concentrations much lower than our initial value (S3 Figure G). *E. coli* does not use overflow metabolism in our model, though *E. cloacea* does, and *E. coli* does when in the presence of glucose (S3 Figure H,I,J).

To test if the direct uptake of oxygen was indeed responsible for *E. coli*’s dominance of the microbiota, we disabled the oxygen uptake reaction of *E. coli* for 30 simulations with 1 µmol initial oxygen per lattice site. Under these conditions, *E. coli* failed to dominate the population (Fig. 5H). Other species, primarily *E. cloacae*, became dominant early in the simulations as a primary consumer (S4 Video). These species are replaced in all simulations by a population composition similar to that from the simulations without oxygen. In some studies, *Klebsiella* species were the dominant early members of the Enterobacteriaceae instead of *E. coli* (50). To examine whether our model could also reproduce early dominance by a *Klebsiella* species we initialized the simulations with a set of species containing *K. pneumoniae* instead of *E. coli*. At 0.1 µmol initial oxygen per lattice site, these simulations behaved the same as the original simulations initialized with *E. coli* (S3 Figure K): *Bifidobacterium* became the most abundant group after 21 days in 26 out of 30 simulations. In simulations initialized with both *K. pneumoniae* and *E. coli, K. pneumoniae* became the most abundant in 15 of 30 simulations, and *Bifidobacterium* in the other 15 (S3 Figure L). Altogether, the model predicts that *E. coli* relies on direct consumption of oxygen to dominate the microbiota under initially oxygenated conditions in the model. It also indicates that *E. cloacea* does not have an anaerobic metabolism competitive enough to sometimes become dominant over *Bifidobacterium* spp. as *E. coli* does, and that other Enterobacteriaceae such as *K. pneumoniae* can perform similarly to *E. coli*.

### Sensitivity analysis

A number of key parameters in our model were based on reasonable estimates. To test the effects of these parameter values on the simulation results, we performed a sensitivity analysis for these parameters under anaerobic conditions, which best represents the situation at the end of the simulations (Fig. 6). As shown in Fig. 2B and S1 Figure D, after relaxing the enzymatic constraint *Bifidobacterium* spp. loses its competitive advantage at high lactose concentrations (S4 Video). This leads to reduced dominance or extinction in whole-gut simulations (Fig. 6A). Tightening the enzymatic constraint led to *Bifidobacterium* spp. remaining dominant, but with a much smaller population size. Thus the presence of an enzymatic constraint sufficiently large for metabolic shifts, but not its exact level, is crucial for the prediction of *Bifidobacterium* dominance. We set the enzyme constraint at a level strong enough to predict the metabolic shifts observed *in vitro*, but not so strong as to prevent reasonable growth. The ATP required for a unit of population growth had a linear effect on population size, as expected, but had little effect on the relative abundance of *Bifidobacterium* (Fig. 6B). The large number of populations with a low ATP requirement also made the model computationally unfeasible. The rate of colonization had little effect on the simulation outcomes: at increased colonization rates, *Bifidobacterium* spp. still dominated the microbiota and its relative abundance remained unchanged (Fig. 6C). Even eliminating colonization altogether hardly affected *Bifidobacterium* spp. dominance in most simulations, but led to complete extinction of all *Bifidobacterium* species in 5/30 simulations and of all 12 non-*Bifidobacterium* species in 9/30 simulations. Thus we kept a moderate amount of colonization in the model. We also examined the effect of placing new populations after initialization only in the first column of the lattice, with a probability increased to match the lower number of eligible squares (S4 Figure A). This also led to a slightly larger but still comparatively small population for the non-*Bifidobacterium* species. To determine the optimal spatial discretization we ran the model on a more coarse and on a more refined lattice, scaling the diffusion, advection, and initialization and colonization parameters accordingly (Fig. 6D; see methods section Population dynamics). Lattice refinement did not change the dominance of *Bifidobacterium* nor the spatial distribution of metabolites and populations (S4 Figure B-E), but on a more coarse lattice the smaller number of lattice sites seemed to increase the rate of extinctions and variability of metabolite distributions (S4 Figure F-I). The diffusion of metabolites had a larger effect on the simulation results (Fig. 6E). At low diffusion rates, the populations tended to die out, because they had less opportunity to take up lactose as they were exposed only as it ‘passed by’ due to advection. At high diffusion rates cross-feeders and *Bifidobacterium* consumed all lactate (S4 Figure J); in infant feces substantial amounts of lactate are found (40, 51). We therefore decided to choose a value for which the system produced the observed concentration of lactate. Finally the amount of mixing of bacteria was varied (Fig. 6F). *Bifidobacterium* remained dominant regardless of the diffusion parameter, even when the bacteria were fully mixed (i.e. placed at random locations) each timestep. Without mixing the populations could not spread, for lack of empty space. The bacterial diffusion constant is 1.3 times higher than the metabolic diffusion constant. As shown in Fig. 6F the exact setting of the bacterial diffusion has little effect on the outcomes of the simulations, so we have maintained bacterial diffusion at this baseline.

**FIG 6.**
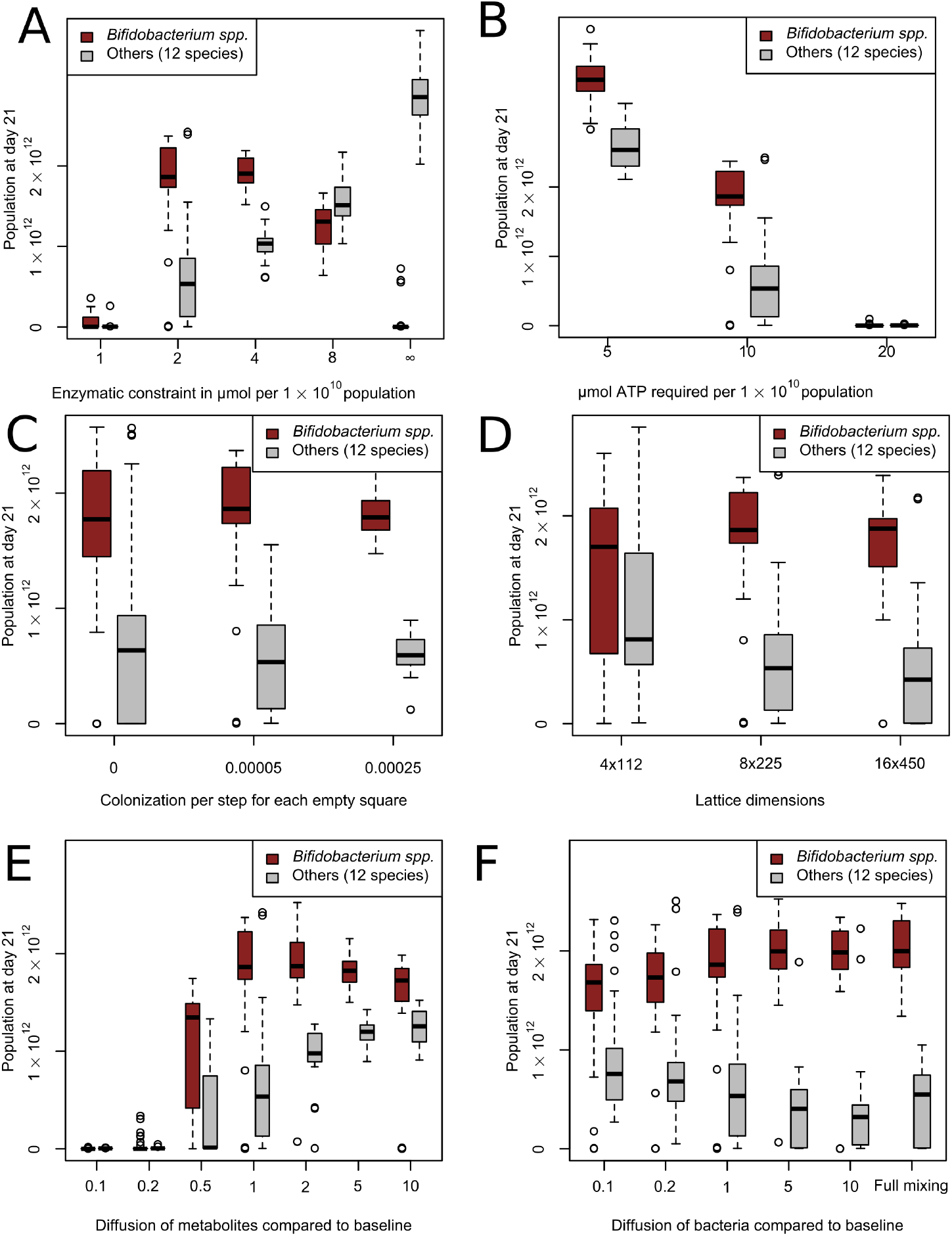
Sensitivity analysis for estimated parameters Abundances for *Bifidobacterium* spp. and other species at the end of day 21 with different parameters. n=30 for each condition. (A) Effect of enzymatic constraint around the default value of 2 µmol flux per timestep per 1·10^10^ population (B) Effect of the amount of ATP required for growth of 1 · 10^10^ bacteria around the default value of 10 µmol (C) Effect of variation in continued influx of populations per timestep per empty square around the default of 0.00005 (D) Effect of variation in width and height of lattice, including adjustment of associated parameters, around the default of 8×225 lattice sites (E) Effect of variation in diffusion for metabolites away from the baseline of 4.4 · 10^−5^ cm^2^/s (F) Effect of variation in diffusion of bacterial populations away from the baseline of 5.7 · 10^−5^ cm^2^/s.

## DISCUSSION

To develop new hypotheses for possible external effects on the initial phases of the infant gut microbiota, we have developed a dynamic multiscale model. Our simulations reproduce the initial dominance of Enterobacteriaceae, particularly *E. coli*, over *Bifi-dobacterium* species, and the subsequent dominance of *Bifidobacterium* species, out of a broad consortium of infant species. Moreover, our simulations suggest show the consistency of the classical hypothesis that this succession could be due to an initial presence of oxygen, followed by microbial depletion (4, 5). These predictions on oxygen suggest that higher levels of initial oxygen may explain the absence or strong delay of *Bifidobacterium* development that is observed in some infants (2). A continuous input of oxygen did not explain the pattern observed *in vivo* (S3 Figure A-D), but stopping the input of oxygen at the midway point of the simulation did lead to a similar pattern (S3 Figure E-F).

While it is known that there is variation in colonic oxygenation (47, 48), the range of this variation in newborns is not known. In premature infants a positive association exists between the number of days of supplemental oxygen and *Klebsiella* sp. abundance (50). *Klebsiella* was the primary representative of the Enterobacteriaceae in this study population. However, only very few of these infants developed a *Bifidobacterium* dominated microbiota, regardless of the duration of supplemental oxygen. Nonetheless, it may be desirable to reduce oxygen concentration faster to achieve the positive health effects of *Bifidobacterium* more consistently. As the direct oxidation of lipids and other organic substrates may contribute to creating anaerobic conditions in the colon (7), altered nutrition may allow for a decrease in oxygen in the colon without stimulating facultative anaerobic bacteria such as *E. coli*. There are also indications that short-chain fatty acids produced by *Bifidobacterium* may decrease oxygen levels by stimulating the oxygen use of colonocytes (8). Future versions of the model could represent these oxygen-depleting mechanisms by localized (e.g., near the intestinal wall) and distributed oxygen consumption terms. At present the model predicts that distributed oxygen usage by such additional processes will speed up the succession to an anaerobic, *Bifidobacterium*-dominated microbiota. The limit case, i.e. extremely fast oxygen consumption, is represented by the anaerobic case (Fig. 3). The effect of localized oxygen consumption processes, e.g., near the intestinal wall is a topic of our ongoing research.

Our simulations predict that the dominance of *Bifidobacterium* spp. in the anaerobic infant system could be explained from its consumption of lactose through its bifid shunt metabolism. Our model can correctly represent bifid shunt metabolism due to the implementation of an enzymatic constraint on the maximum total metabolic flux in the bacteria (Fig. 2). Both the low and high yield metabolism, and the switch between them based on nutrient availability, have been extensively described *in vitro* (38). The FBA with enzymatic constraint method we use is similar to flux balance analysis with molecular crowding (FBAwMC) (21). As in FBAwMC we consider the enzymatic constraint to represent the limited availability of space and production capacity for enzymes within a bacterial cell. However, in the absence of good reaction-specific thermodynamic information for our current model, we do not utilize the reaction-specific technique of FBAwMC. We instead base an enzymatic constraint only on the size of the local population, which is sufficient for modeling a metabolic switch. This is equivalent to using FBAwMC and placing all crowding coefficients at the same positive number, which has been done previously to model metabolic switches (21). More broadly, our method is similar to other existing methods that model metabolic switches by including limits on fluxes that represent some physical limit on metabolism, such as proteome constraints or membrane occupancy (52, 35, 53, 54). It has been shown that metabolic switches in microbes can be modelled in many ways, provided that two simultaneous constraints are in place (22). In our case, these are the concentration of substrate and the enzymatic constraint.

The metabolic switch of *Bifidobacterium* in our model determines the production of its metabolites. The two most abundant, acetate and lactate, are present in the simulated feces at a median ratio of 3.5:1 (Fig. 3B), similar to that found in formula-fed infant feces (40, 51). We can derive a concentration in mM if we assume each of the eight lattice sites whose metabolites are advected out of the system each step contains 0.05ml of water. This is based on an estimated total volume of 90ml, divided over 1800 lattice sites (Table 3). This results in an average concentration of 29 mM acetate and 8 mM lactate in the simulated fecal output analyzed in Fig. 3B. These values are close to, and well within the variability, of the average values for acetate and lactate of 31 and 12 mM respectively in the feces of two-week old infants in (51). The median ethanol:acetate ratio of 1:21 is similar to the ratios of 1:18 and 1:10 reported in term infants (43) and the 1:50 reported in pre-term infants (55). We use ratios here because the measures per gram dry weight are not available in our model. We observe a lower quantity of ethanol in the simulated fecal output of the simulations where lactate uptake by *Bifidobacterium* was disabled (S2 Figure E).

**TABLE 3.** Default parameters of the model

| Parameter | Value | Unit |
| --- | --- | --- |
| Timestep | 180 | seconds |
| Lattice site side length ($\Delta x$) | 2 | millimetre |
| Width of the lattice | 225 | lattice sites |
| Height of the lattice | 8 | lattice sites |
| Colon transit time | 11 | hours |
| Average number of initial populations | 540 | - |
| Initial size per population | $5 \cdot 10^7$ | number of bacteria |
| Population size required for division | $1 \cdot 10^{10}$ | number of bacteria |
| Death probability | 0.0075 | per timestep per population |
| Enzymatic constraint | 2 | $\mu\text{mol}$ flux per timestep per $1 \cdot 10^{10}$ population |
| New species placement probability | 0.00005 | per timestep per empty lattice site |
| Diffusion of metabolites | $4.4 \cdot 10^{-5}$ | $\text{cm}^2/\text{s}$ |
| Diffusion of populations | $5.7 \cdot 10^{-5}$ | $\text{cm}^2/\text{s}$ |
| Growth per $\mu\text{mol}$ ATP | $1 \cdot 10^9$ | number of bacteria |
| Lactose input | 211 | $\mu\text{mol}$ per three hours |

*Bifidobacterium* takes up lactate in our model and converts it into pyruvate, catalyzed by lactate dehydrogenase. In our model this pyruvate is then further converted to formate, ethanol, and acetate through the high-yield pathway also used in lactose metabolism. However, strong lactate uptake by *Bifidobacterium* spp. has not been observed *in vitro* (in e.g. (56)). Thus, while the enzymes used for lactate consumption exist in *Bifidobacterium*, it is not known if the pathway is used *in vivo* to the extent predicted by the model. Lactate uptake by *Bifidobacterium* has only little impact on the simulation results (S2 Figure D). Similarly, though growth on lactate by *E. coli* in the presence of oxygen is well documented, this is not the case for the anaerobic condition (57). Lactate uptake is not essential for *E. coli*, as it can exist in a more purely primary consumer role (Fig. 4F). The third consumer of lactate, *B. hansenii*, is present only marginally *in vivo* (11). This matches with our results for initially oxygenated conditions, in which *B. hansenii* is also largely absent. However, other species that could fill the same lactate-consuming secondary consumer niche as *B. hansenii* in our model are common *in vivo* (58). Our selection of species may be further improved to allow the correct species to emerge in this secondary consumer niche. For example, it might be appropriate to add *Eubacterium hallii*, a common infant gut bacterium known to consume *Bifidobacterium*-produced lactate and acetate *in vitro* (59).

More generally, our model predicts a lower diversity overall in both bacterial species and metabolites compared to what is observed *in vivo* (11). While the abundant species in our model largely match with the abundant species *in vivo*, many other species such as those of the genera *Streptococcus* and *Lactobacillus* have a lower abundance in our model compared to the *in vivo* conditions(11). We have decided to keep all less abundant species in our simulations, to show how in most cases the correct species out of a broad consortium become dominant. There were no large differences when we ran the simulation without these smaller species (S2 Figure A). Particularly the lack of representation of the Bacilli species in our model outcomes is notable. While some sources report a portion (e.g. 12-14% (2)) of their subjects to be dominated by Bacilli, mainly *Streptococcus*, the *Streptococcus* species in our model never became dominant. We showed that *S. salivarius* has an inferior metabolism on lactose and lactate compared to other species in our model (Fig. 2A, Fig. 4E, Fig. 5G).

The discrepancy between our model and the *in vivo* data in the abundance of less common species such as *Streptococcus* may partially be explained through the focus of our model on carbon dissimilation. Though it would be preferable to represent the whole metabolism of the infant microbiota, including factors such as consumption of amino acids, oligosaccharides and intestinal mucus, the initial version presented here only considers carbon metabolism from lactose. While a more extensive metabolism might have allowed more mechanisms and niches to be discerned, it would also introduce additional free parameters, as there is no clear data on the concentration and uptake of these nutrients. The uptake bounds would have to be set arbitrarily, with uniform values or random sampling. Many substances such as fatty acids or protein residues are not even included as metabolites in the database we use for bacterial metabolism (23), further complicating their introduction to the model. By focusing on carbon metabolism, using ATP production as a proxy for growth rate, and only using lactose as an input nutrient, we can circumvent these problems. ATP production has been shown to be a good proxy for biomass production in *E. coli* (31). Supplementation of the *in vitro* and model organism infant gut microbiota with prebiotic carbohydrates led to a larger microbiota, primarily due to larger *Bifidobacterium* populations(24, 25). This indicates that carbon metabolism is a limiting factor, especially in the absence of prebiotics, as in our model. However, as the increased population in these *in vitro* and *in vivo* studies consisted largely of *Bifidobacterium*, other species were not necessarily also carbon limited. In addition, ATP production may not be a good proxy for biomass production in some species. Species-specific information on the relation between ATP and biomass production may aid future modelling. Finally, gaps in the GEMs we use may cause ATP production itself to be underestimated compared to what occurs *in vivo*.

Besides details on the modelling of metabolism, the way that diffusion and advection of bacteria and metabolites is handled in the simulations may impact the predictions of our model. The sensitivity analysis demonstrated an unrealistic metabolic output when the metabolic diffusion coefficient was raised to 2.2 · 10^−4^ cm^2^/s (S4 Figure J). Although this value is much lower than diffusion coefficients measured in the adult intestinal lumen (1 · 10^−2^ cm^2^/s) (60), in the infant colon mixing may be reduced compared to the adult colon: motor activity that mixes the colon contents (68) is rarer in infants than in older children or adults (61, 62). Exact measurements of mixing in the infant colon will be hard to obtain. Related to this, while we did not examine the effect of bacterial advection in our current work, our previous work (18) suggested that increased bacterial advection greatly reduces diversity and spatial patterning in the model. In fact, compared to the adult gut, in the infant gut there is increased motor activity driving advection (62) and infants display faster colonic transit than adults (32, 33). In light of these data, which seem to indicate much faster bacterial advection and stronger luminal mixing than what we have assumed in our model, a potential interpretation of the present set-up of our model is that it represents the dynamics of bacterial populations adhering to the intestinal wall in interaction with metabolites advecting through the lumen. Our future work will explore in more detail how a balance between adhesion of microbiota to the mucus (63), versus advection of bacteria and metabolites in the lumen affects the colonization of the infant colon.

Our modeling approach relates to alternative simulation frameworks, such as Steadycom, BacArena, and COMETS (16, 64, 17, 65). These frameworks, in particular the new COMETS 2 framework (19) would certainly be suitable for answering questions similar to those asked in the present work. In absence of suitable frameworks at the initial phases of this project, and given the flexibility that comes with using an in-house code base, we preferred to continue developing our own line of gut models (18). A future implementation of our model in one of the available simulation frameworks would be useful exercise, facilitating future development and comparison of models.

We have used our modeling approach to generate testable hypotheses on the causes and mechanics of succession in the infant gut microbiota, potentially laying the foundation for nutritional interventions that could improve the health of infants. It should be emphasized that the current and future work on this model represent a tool for generating hypotheses, and for testing potential mechanical explanations. We cannot fully represent the complexities of the infant gut microbiota, and any generated hypotheses must necessarily be validated *in vitro* and *in vivo*. There is work to be done to further study the relevant factors, and to bring these results into a more realistic model context. We aim to do so by integrating the prebiotic oligosaccharide and protein content of nutrition into the model. This may lead to the creation of more niches in the model, and so a diversity closer to that of the *in vivo* system. We will also continue to improve the selection and curation of metabolic models, such as by disabling lactate uptake in certain species or including a wider diversity of GEMs. This may lead to novel future insights on the interactions between differences in infant nutrition and succession in the infant microbiota.

## METHODS

We use a spatially explicit model to represent the newborn infant microbiota (Fig. 1A). Our model is based on an earlier model of the adult gut microbiota (18). The model consists of a regular square lattice of 225 × 8 lattice sites, where each lattice site represents 2 *mm* × 2 *mm* of space, resulting in an infant colon of 450mm by 16mm. Each lattice site can contain any number of metabolites of the 723 types represented in the model in any concentration and a single simulated bacterial population. The metabolism of these bacterial populations is based on genome-scale metabolic models (GEMs) from the AGORA collection of species-specific GEMs (23). From this set, we have chosen 15 GEMs based on a consortium of known infant microbiota species (30, 11). Their metabolic inputs and outputs are calculated using dynamic flux balance analysis (FBA) (66) with an enzymatic constraint functioning as a limit on the total flux through the network (20). The effects of the FBA solution are applied to the spatial environment and then recalculated each timestep, creating a spatial dynamic FBA.

We identify the two narrow ends of the rectangle with the proximal and distal ends of the colon. In each timestep metabolites both mix and flow from the proximal to the distal end. Bacterial populations are mixed, but do not flow distally as metabolites do. At the start of each run, we initialize the system with a large number of small populations. We let these perform their metabolism each timestep. They take metabolites from the environment, and deposit the resulting products. We let the populations grow and divide according to their energy output. Both the initial placement locations and movement of the populations is random, introducing stochasticity in the model.

The system develops differently depending on initial conditions, into a diverse and complex ecosystem.

### Species composition

Each population is represented by a GEM of a species. 15 different metabolic models are used in our spatially explicit model (Table 1), selected based on previous research (30). From this list of genera, the most prevalent species within that genus in the vaginally born newborn data set from Bäckhed *et al*. was used (11). One group from the list could only be determined at the family level - Lachnospiraceae. *Ruminococcus gnavus* was chosen to represent this group, based on its high prevalence among this group and the prior inclusion of species from the *Blautia* and *Dorea* genera (69). Because the genus *Bifidobacterium* is known to be particularly diverse (70), we represented it with models of three different strains. All models are based on the GEMs created by Magnúsdóttir et al. in the AGORA collection (23).

### Changes from AGORA

We use updated versions of the AGORA GEMs (74), to which we have applied checks and modifications. Firstly, the objective function was changed from the biomass reaction included in the models to a reaction only requiring ATP production. As this reaction yields only ADP and P_i_, it is mass-neutral. This allows us to focus on carbon dissimilation within the GEMs, and the differences in underlying metabolism. ATP yield has been a good proxy for biomass production in previous studies (75). Focusing on carbon dissimilation means we can leave all unknown uptake bounds at 0, instead of using an arbitrary or randomized level, as in some other studies (17, 16).

We also checked the metabolic networks for reactions that allow for the occurrence of unrealistic FBA solutions, and we have added additional reactions. In the *Bifidobacterium* models the ATP synthase reaction was made reversible. This allows it to function as a proton pump, which *Bifidobacterium* species use to maintain their internal pH (76). Lactose permease reactions were added to all *Bifidobacterium* species in the model (77) based on those available in other models in the AGORA collection (23). Combined with the existing reactions in the model and the metabolic constraints this leads to a set of reactions simulating a realistic bifid shunt yield, when substrate is abundant, of 5 mol ATP, 2 mol lactate and 3 mol acetate per mol of lactose(38, 78). Lactose permease was also added to *Streptococcus salivarius* and *Ruminococcus gnavus* to bring them in line with existing literature on their *in vitro* behavior (79, 80). *Veillonella dispar* is the only species in the model that does not have any lactose uptake (71). A complete list of changes is presented in S1 Table.

### Checking the validity of the GEMs

After the changes in S1 Table were applied all GEMs used in the model were tested individually to ensure that they could grow on a substrate of lactose. Only *Veillonella* did not pass this test, which is consistent with *in vitro* observations (71). *Veillonella* did pass when lactate, a common infant gut metabolite, was used instead. We also tested all GEMs individually for spurious growth in absence of substrates. To this end, all uptake bounds except water were set to 0. None of the GEMs grew under these conditions. All models were tested for having a net neutral exchange of hydrogen, carbon, oxygen and nitrogen at each timestep of the model during the simulations. There should be no net change in the number of atoms in the medium due to the calculated fluxes, because the reaction we set as an objective function does not remove any atoms, and there are no other sinks in the simulations. In the present simulation, only water, oxygen, and lactose are introduced into the simulation, so no other atoms than these three are considered. The tests revealed two errors in glycogen metabolism in several GEMs in AGORA that resulted in a energetically-favorable removal of intracellular protons from the system. We corrected these GEMs by replacing the reactions responsible for this erroneous energy source (S1 Table). With the corrections we applied to the model, these tests always passed - allowing for rounding errors of less than 1 · 10^−8^ µmol per FBA solution.

The thermodynamic correctness of all reactions was checked by calculating the net difference in Gibbs free energy between input and output metabolites, using a pre-existing dataset of thermodynamic source data (72, 73). The conditions assumed for the calculation of the Gibbs free energy were a pH of 7 and an ionic strength of 0.1M (72). This was recorded for each population at each timestep over the course of a full run of 10080 timesteps. 99.999% of all FBA solutions in the simulations of Fig. 3 passed the test by containing less free energy than the associated inputs. The remaining 0.0001% all had an amount of energy in the outputs equal to that of the inputs. All of these solutions had very low growth rates, less than 0.001% of the average growth rate. The sum of these growth rates over 30 simulations was less than 0.05 of an initial Spopulation, totalling 2 · 10^6^ bacteria.

### FBA with enzymatic constraint

Each timestep of the model, a modified version of flux balance analysis with an enzymatic constraint is used by each population to determine what reactions it should use to achieve the most biomass production from the metabolites available to it (81, 20). First, each GEM is converted to a stoichiometric matrix *S*. All reversible reactions are converted to two irreversible reactions, so that flux is always greater than or equal to 0. All reactions identified as exchange, sink, or demand in the metabolic reconstruction are marked as exchange in the matrix. These reactions exchange nutrients or metabolites with the environment. Each timestep, all reactions are assumed to be in internal steady state (81):

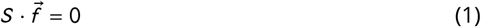

Where 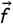 is a vector of the metabolic fluxes through each reaction in the network (in mol per time unit per population unit).

Each exchange reaction that takes up metabolites from the environment (*F*_*i n*_ (*m*)) is constrained by an upper bound *F*_*ub*_ (*m*) which represents the limited availability of most nutrients from the environment:

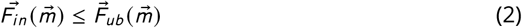

Where 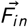 is a vector of fluxes between the environment and the bacterial population (in mol per time unit per population unit) and 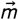 is a vector of all metabolites. 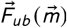 is set dynamically at each timestep *t* by the spatial environment (discussed in detail below) at each lattice site *x* :

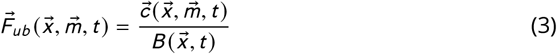

Where 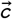 is a vector of all metabolite concentrations in mol per lattice site, 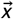 is a vector of all lattice sites and 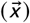 is the size of the local bacterial population in population units. Population units are continuous, the size of *B* can range from 5 · 10^7^ to 2 · 10^10^, or 4 · 10^10^ when division is not possible.

There is an additional constraint on the total flux. This constraint represents the limited amount of metabolism that can be performed per cell in each timestep. Each cell can only contain a limited number of enzymes, and each enzyme can only perform a limited number of reactions in limited time. Not limiting flux would lead to an unrealistic situation. The enzymatic constraint uses the sum of all fluxes 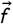, not just those that exchange with the environment:

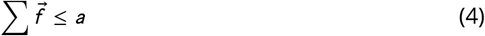

The enzymatic constraint *a* is in mol per time unit per population unit (Table 3). As both 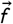 and *a* are per population unit, this limit scales with population size, allowing each bacterium to contribute equally and independently to the metabolic flux attained in a lattice site. The enzymatic constraint is included as a constraint on each local FBA solution separately at each timestep. The enzymatic constraint was first proposed in a metabolic modelling context in 1990 (20), adapted from the study of other capacitated flow networks. In our context the enzymatic constraint represents the limited availability of physical space for enzymes. Our method is similar to FBA with molecular crowding in that we use an additional constraint to model metabolic capacity (21). However, we do not utilize the necessarily reaction-specific crowding coefficients.

FBA then identifies the solution space that adheres to these constraints and from this spaces identifies the solution that optimizes the objective function. The set of solutions consists of a set of exchange fluxes 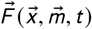 and a growth rate 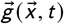. These exchange fluxes are taken as the derivatives of a set of partial-differential equations to model the transport of intermediary metabolites (see below). The size of the population increases in proportion to the growth rate produced by the solution.

### Environmental metabolites

We modeled a set of 723 extracellular metabolites - the union of all metabolites that can be exchanged with the environment by at least one GEM in the model. In practice, only a fraction of the metabolites occur in any exchange reaction that has flux over it, and only 17 metabolites are ever present in the medium in more than micromolar amounts in our simulations outside of the sensitivity analysis (Table 2). Though the model distinguishes between L-lactate and D-lactate, we display them together in our figures. Nearly all lactate produced and consumed in our model is L-lactate.

To mimic advection, the complete metabolic contents of the lattice except oxygen are moved towards the distal end by 2mm (one lattice site with the default parameters) per timestep, i.e. once every 3 simulated minutes. This leads to an average transit time of approximately 11 hours, in agreement with the observed cecum to rectum transit time in newborns (32, 33). Metabolites moving out of the distal end are removed entirely and analysed separately (see Section ‘Analysis’ for detail).

Every timestep all metabolites diffuse to the four nearest neighbours 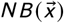 at an equal rate for all metabolites (Table 3). We used a baseline of 4.4 · 10^−5^ cm^2^/s to represent mixing of metabolites, which is an order of magnitude higher than normal diffusion for common metabolites (82) to represent active mixing due to colonic contractions. Metabolites are also added and removed by bacterial populations as a result of the FBA solutions, yielding

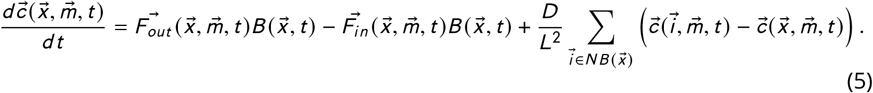

Where *F*_*out*_ is a vector of fluxes from the bacterial populations to the environment, in mol per time unit per population unit.

All lattice sites initially contain water as their only metabolite, except for the conditions where oxygen is added. Metabolites representing the food intake are inserted into the first six columns of lattice sites every three hours (60 timesteps) to approximate a realistic interval for neonates (83). In the current model this food intake consists solely of lactose, in a concentration in line with human milk (84), assuming 98% host uptake of carbohydrates before reaching the colon, a commonly used assumption (16). Water is provided as a metabolite in unlimited quantities. Oxygen is placed evenly distributed or at the upper and lower boundaries in some simulations. No other metabolites are available, other than those produced as a result of bacterial metabolism within the model.

### Population dynamics

Each local population solves the FBA problem based on its own GEM, an enzymatic constraint *a*, its current population size 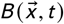 and the local concentrations of metabolites 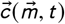 at each timestep, and applies the outcome to the environment (see above) and its own population size, as follows:

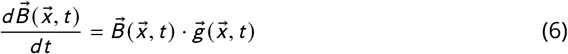

Populations at least 200 times the initial size (Table 3) will create a new population in one empty adjacent lattice site, if possible. Half of the old population size is transferred to the new population, in such a way that the total size is preserved. To mimic colonization events new populations are introduced at random into empty lattice sites during the simulation, representing both dormant bacteria from intestinal crypts (34) and small bacterial populations that are formed from ingested bacteria, which may only become active after having diffused far into the gut. Each empty lattice has a probability of 0.00005 (Table 3) each step to acquire a new population of a randomly selected species, as follows:

**for** *site* ∈ *latticesites* **do**

**if** *site* == *empt y* **then**

**if** *randomnumber* ≤ *placement probability* **then**

Place metabolic network of random species at *site*

The probability is scaled by 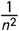, with *n* the scaling factor of Δ*x*, the side length of a lattice site. There is an equal probability for any species in the model to be selected. As we consider these new populations to be new colonizers we initialize them at the same population size *B* as the initial populations in the model (Table 3). Each population dies out at a probability of 0.0075 per timestep, creating a turnover within the range of estimated microbial turnover rates in the mouse microbiota (85).

To mimic the mixing of bacterial populations, the lattice sites swap population contents each timestep. We use the following algorithm, inspired by Kawasaki dynamics (86), as also used previously for bacterial mixing (18, 87): In random order, the content of each site, i.e., the bacterial population represented by its size and the GEM but not the metabolites, are swapped with a site randomly selected from the set consisting of the site itself and the first and second order neighbours. This swap only occurs if both the origin and destination site have not already swapped. Bacterial populations at the most distal column, i.e. at the exit of the colon, are deleted from the system. With this mixing method the diffusion constant of the bacterial populations becomes 5.7 · 10^−5^ cm^2^/s (Table 3). In the simulations that use a finer or a coarser grid (Fig. 6D) the number of swaps is scaled as 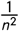, with *n* the scaling factor of Δ*x*, the side length of a lattice site, thus maintaining the same diffusion rate. For *n* < 1 all sites are marked as unswapped once all sites have attempted a swap. This allows for sites to swap multiple times. The same approach was used to change the bacterial diffusion rate in the sensitivity analysis (Fig. 6F). To achieve full mixing all bacteria were assigned random non-overlapping locations at every timestep. Note that the diffusion rate of bacterial populations with the default parameters (Table 3) is 1.3 times higher than that of the metabolites. As shown by the sensitivity analysis (Fig. 6F) this has little effect on the results.

### Initial conditions

The simulation is initialized by placing a number of very small populations (*B*) of the various species randomly across different lattice sites of the environment (Table 3). There is a probability of 0.3 for each lattice site to acquire a population, an average of 540 for our lattice. The size of initial populations is scaled to be roughly equivalent to a plausible initial total load of approximately 3 · 10^10^(88), assuming a total colon volume of approximately 90 ml. As there is little information on the relative abundance of species in the very early infant gut, we place all species with equal probability. In initially oxygenated conditions, oxygen is also placed as a metabolite now. Water is always considered to be present everywhere. No other metabolites are initially present except for the first feeding.

The ratio of population to bacterial abundance was chosen as a balance between a high bacterial resolution and computational complexity, as the FBA solution of each population is calculated independently per timestep. This resolution splits the infant microbiota into approximately 1-600 units (depending on conditions), allowing for the presence of species at relatively low abundances, and spatial variation. There is no hard limit on the number of populations in the model - it is limited only by the balance between growth (through consumption) and death.

## Analysis

At each timestep we record the location and exchange fluxes 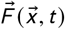 of major metabolites as well as the size 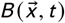 and species of all populations. This is used to analyze both population composition and metabolic fluxes over time and space. In addition, each timestep we record the location and quantity 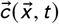 of all metabolites present in more than micromolar quantities. We also record this metabolite data for all metabolites exiting the system at the distal side. This is considered the closest equivalent to a fecal composition in the model, and these results are compared to data from *in vivo* fecal samples.

To detect any irregularities, we also record the net carbon, hydrogen, and oxygen flux of every population and the system as a whole. The difference in Gibbs free energy per timestep is also recorded per for each population per FBA solution, and seperately over the whole system. Estimated Gibbs free energy is derived from the Equillibrator database (73). Energy loss *l* in joules per timestep per population unit is recorded as follows, where *m* are metabolites, *F* is the exchange fluxes in mol per population unit, *B* is the population size and *E* contains the Gibbs free energy in joules per mol for each metabolite,

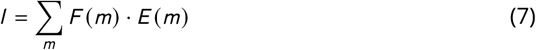

For specific simulations, reactions are removed from the models. This is performed by deleting some reactions from the GEM before the conversion to the stoichiometric matrix. To remove Fructose-6-phosphate phosphoketolase, we removed the reactions R_PKL and R_F6PE4PL. For other simulations, the uptake of certain metabolites is disabled. This is done by placing the upper flux bound 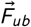 of the relevant exchange reaction at 0 for the relevant populations.

### Parameters

Relevant parameters are listed in Table 3. Based on measurements of the typical length and diameter of the infant colon (26, 27) we estimated a volume of 90 ml. Combined with the average abundance per ml of around 10^10^ after the first days (88), this leads to a very rough estimate of 10^12^ bacteria in the young infant colon. To remain computationally feasible, while still modelling at a high resolution, we divide this population into units of at most 1 · 10^10^. Local populations at or above this limit will divide into two equally sized populations when space is available. This prevents the local population from becoming unrealistically high. Local populations of more than 2 · 10^10^, which can only form if no space is available to divide for a longer time period, cease metabolism.

The lactose input is estimated from the known intake of milk, its lactose concentration, and an estimate of pre-colonic lactose absorption of 98% (89, 16). Little data is available on the growth rate of bacteria within the human colon. Growth rates are expected to be much lower than those found in *in vitro* cultures of individual species (90). In absence of precise data for infants, here we use a death probability that places the replacement rate within the range of estimated doubling times of the whole gut microbiota in mice (85). The colonic transit time is based on data for total transit time gathered with carmine red dye (32), adjusted for the mouth to cecum transit time (33). The timestep interval was set at three minutes, to be able to capture individual feedings at a high resolution. Other parameters were selected to reach plausible outcomes on a metabolic and species level while maintaining computational feasibility. We do not expect the parameters to match precisely with *in vivo* values, if these were measured.

### Implementation

The model was implemented in C++11. Our code was based on code used earlier to model the gut microbiota (18). Random numbers are generated with Knuth’s subtractive random number generator algorithm. Diffusion of metabolites was implemented using the Forward Euler method. The GEMs are loaded using libSBML 5.18.0 for C++. We used the GNU Linear Programming Kit 4.65 (GLPK) as a linear programming tool to solve each FBA with enzymatic constraint problem. We used the May 2019 update of AGORA, the latest at time of writing, from the Virtual Metabolic Human Project website (vmh.life). Python 3.6 was used to extract thermodynamic data from the eQuilibrator API (December 2018 update) (73) and determine mean square displacement of our bacterial diffusion. Model screenshots are made using the libpng16 and pngwriter libraries. Other visualisations and statistical analysis were performed with R 4.1.2 and Google Sheets.

## Data availability

The code used for the model is available from GitHub at https://github.com/DMvers/IGMOST. The data files used for the model are available from GitHub at https://github.com/DMvers/IGMOSTdatafiles.

## SUPPLEMENTAL MATERIAL

**S1 Figure.**
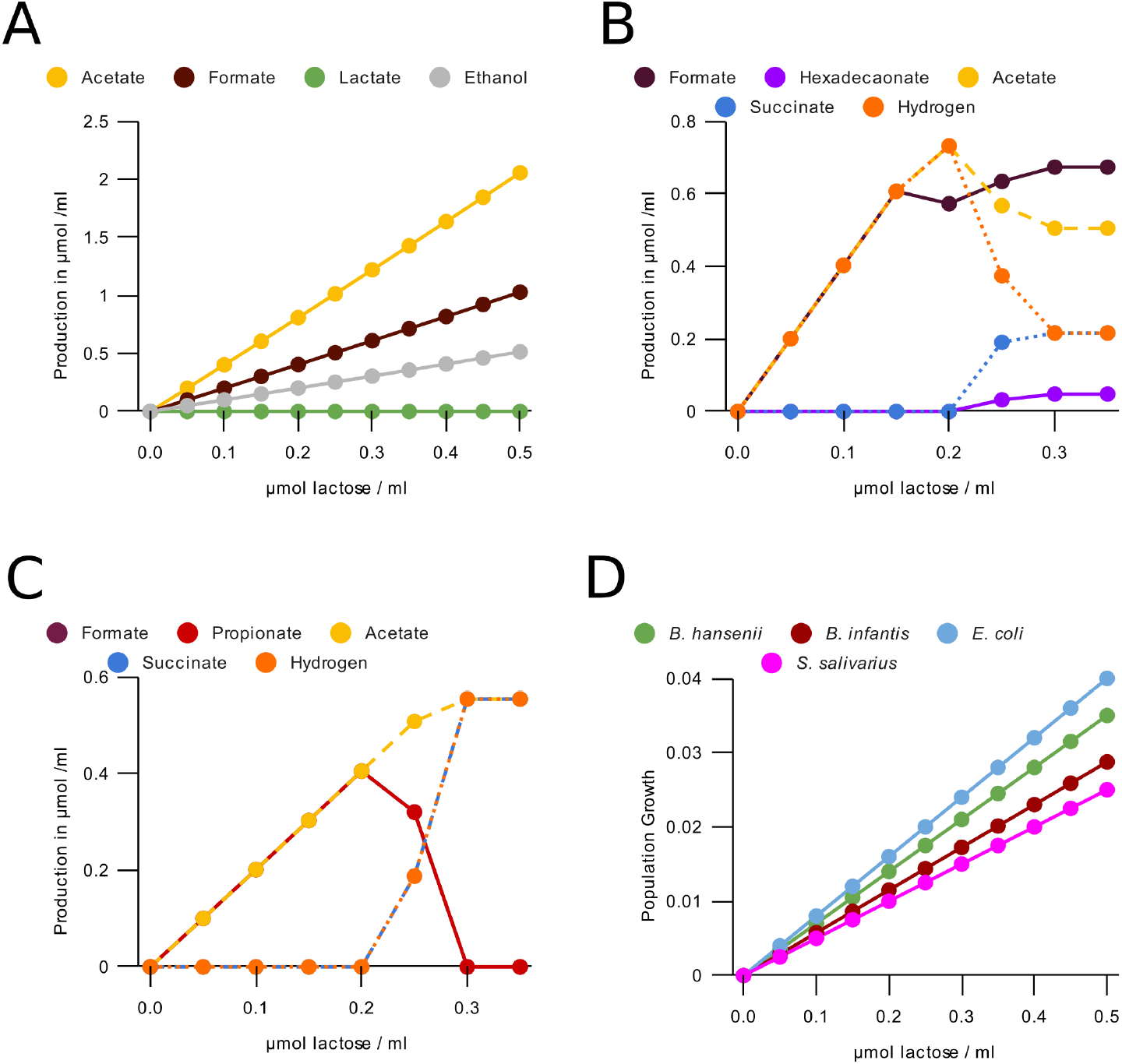
(A) Growth per timestep by lactose concentration for populations of 5 · 10^9^ bacteria with access to one lattice site (0.05ml) of some major bacterial species with the enzymatic constraint disabled (B) Production of metabolites per timestep by an *E. coli* population of 5 · 10^9^ bacteria with access to one lattice site (0.05ml) (C) Production of metabolites by a *B. vulgatus* population of 5·10^9^ bacteria with access to one lattice site (0.05ml). Formate and propionate formation are always equal, and so are hydrogen and succinate. (D) Growth per timestep by lactose concentration for populations of 5·10^9^ bacteria with access to one lattice site (0.05ml) of some major bacterial species with the enzymatic constraint disabled.

**S2 Figure.**
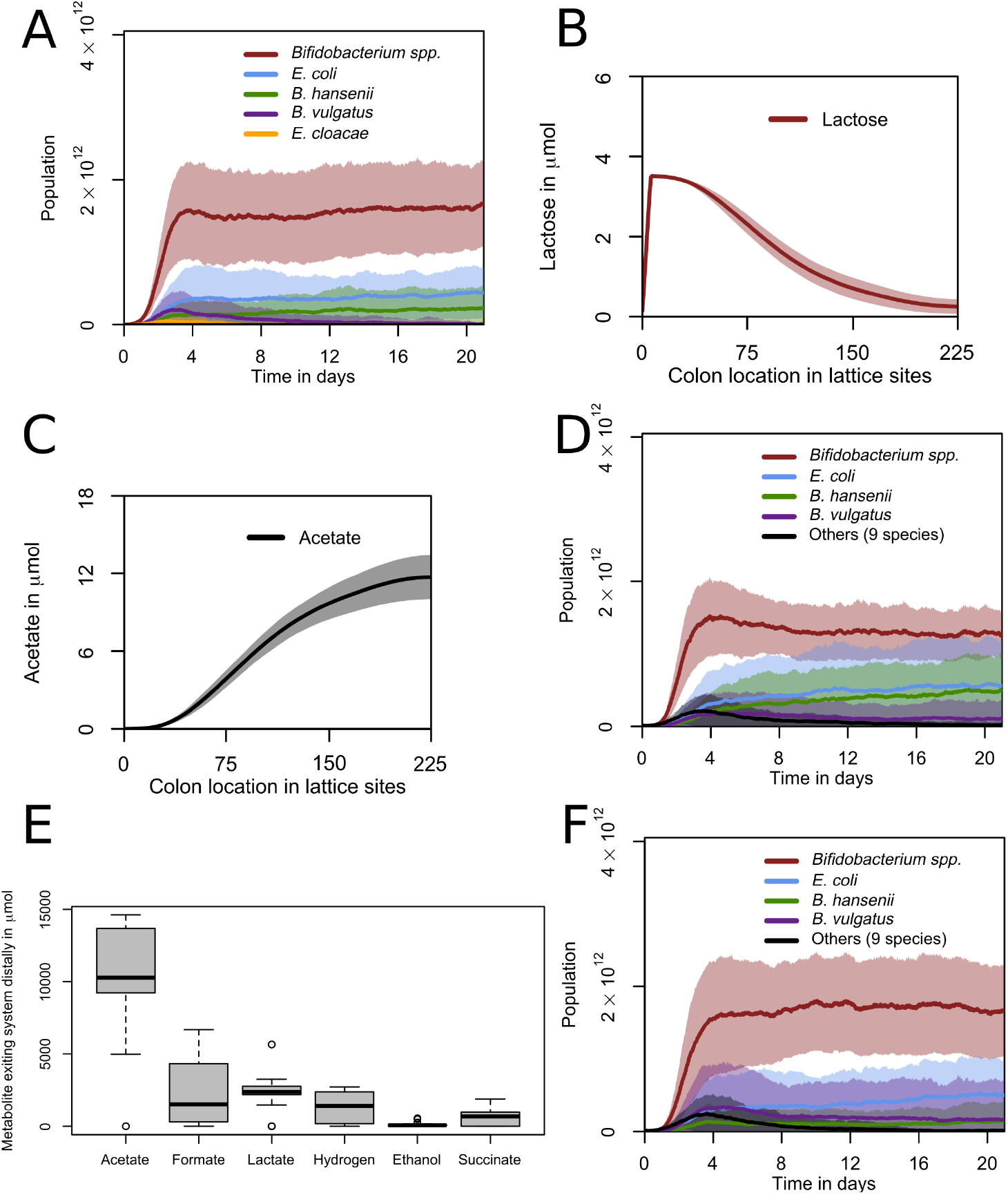
(A) Populations over time with bacteria in the “others” category of Fig. 5 left out. Abundances for grouped species over 21 days. One day is 480 timesteps. Curve shows mean value and shaded area one standard deviation over n=30 simulations. (B,C) Spatial distribution of (B) lactose and (C) acetate over the last two days (960 timesteps) of the simulation. Curve shows mean value and shaded area one standard deviation over n=30 simulations. (D) Populations over time with lactate uptake disabled for *Bifidobacterium* species. Abundances for grouped species over 21 days. One day is 480 timesteps. Curve shows mean value and shaded area one standard deviation over n=30 simulations. (E) Distribution of metabolites exiting the system distally over the last two days (960 timesteps) of the simulation in D. n=30. (F) Populations over time with lactate production disabled for *Bifidobacterium* species. Abundances for grouped species over 21 days. One day is 480 timesteps. Curve shows mean value and shaded area one standard deviation over n=30 simulations.

**S3 Figure.**
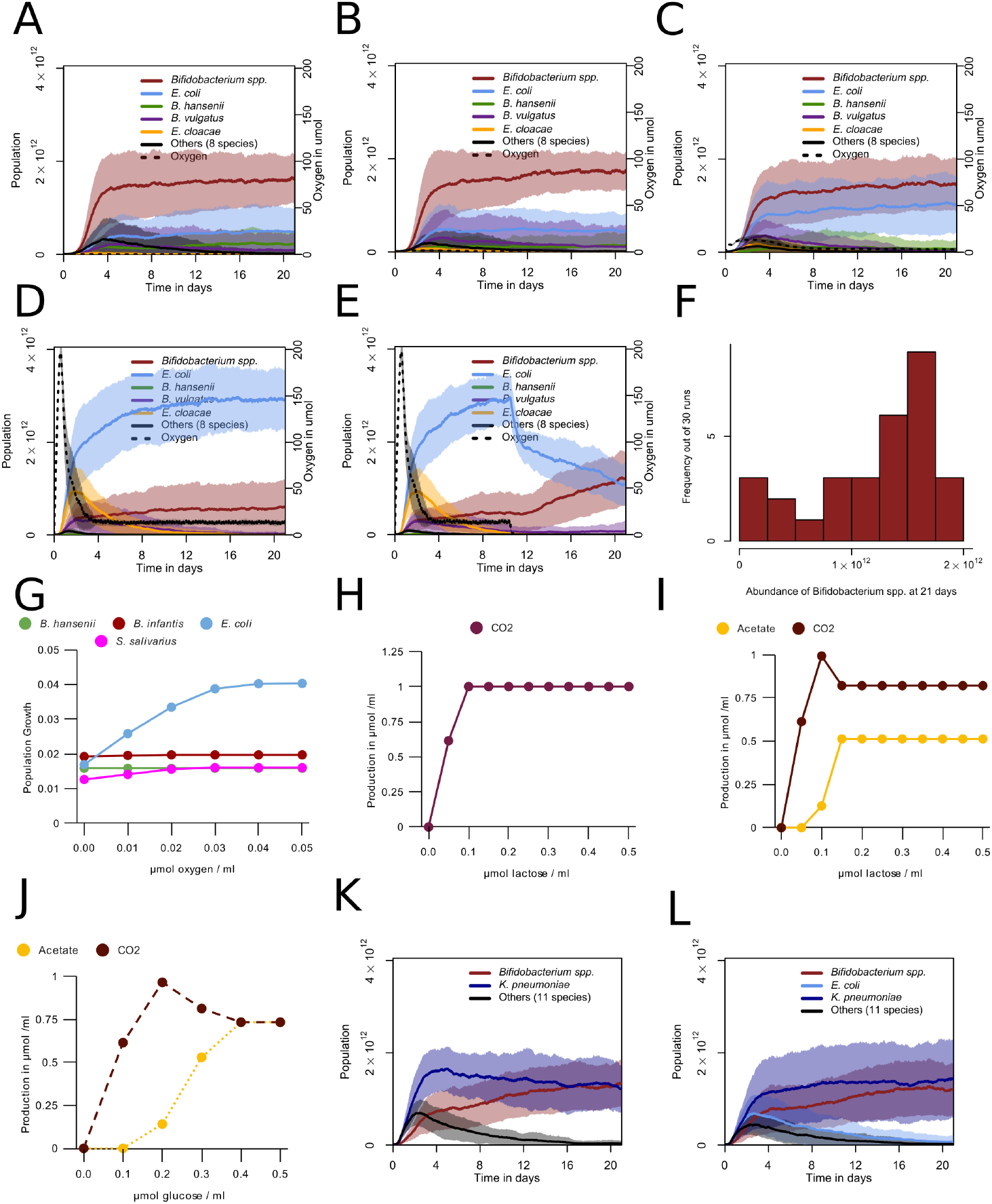
(A,B,C,D) Populations over time with (A) 0.001 (B) 0.01 (C) 0.1 (D) 1 µmol oxygen released in total per step divided over the top and bottom rows. Abundances for grouped species over 21 days. One day is 480 timesteps. Curve shows mean value and shaded area one standard deviation over n=30 simulations per condition. (E) Populations over time with 1 µmol oxygen released per step initially (as in D), decreasing to 0 at step 5040. Abundances for grouped species over 21 days. One day is 480 timesteps. Curve shows mean value and shaded area one standard deviation over n=30 simulations per condition. (F) Distribution of total *Bifidobacterium* abundance at 21 days (10080 timesteps) corresponding to the simulations of E (G) Growth on unlimited lactose per timestep by oxygen concentration for populations of 5 · 10^9^ bacteria with access to one lattice site (0.05ml) of some major bacterial species (H) Production of metabolites by lactose concentration in the presence of abundant oxygen for an *E. coli* population of 5 · 10^9^ bacteria with access to one lattice site (0.05ml) (I) Production of metabolites by lactose concentration in the presence of abundant oxygen for an *E. cloacea* population of 5 · 10^9^ bacteria with access to one lattice site (0.05ml) (J) Production of metabolites by glucose concentration in the presence of abundant oxygen for an *E. coli* population of 5 · 10^9^ bacteria with access to one lattice site (0.05ml) (K) Populations over time with *K. pneumoniae* instead of *E. coli* and 0.1 µmol initial oxygen per lattice site. Abundances for grouped species. One day is 480 timesteps. Curve shows mean value and shaded area one standard deviation over n=30 simulations. (L) Populations over time with *K. pneumoniae* in addition to the 15 species from table 1, and 0.1 µmol initial oxygen per lattice site. Abundances for grouped species. One day is 480 timesteps. Curve shows mean value and shaded area one standard deviation over n=30 simulations.

**S4 Figure.**
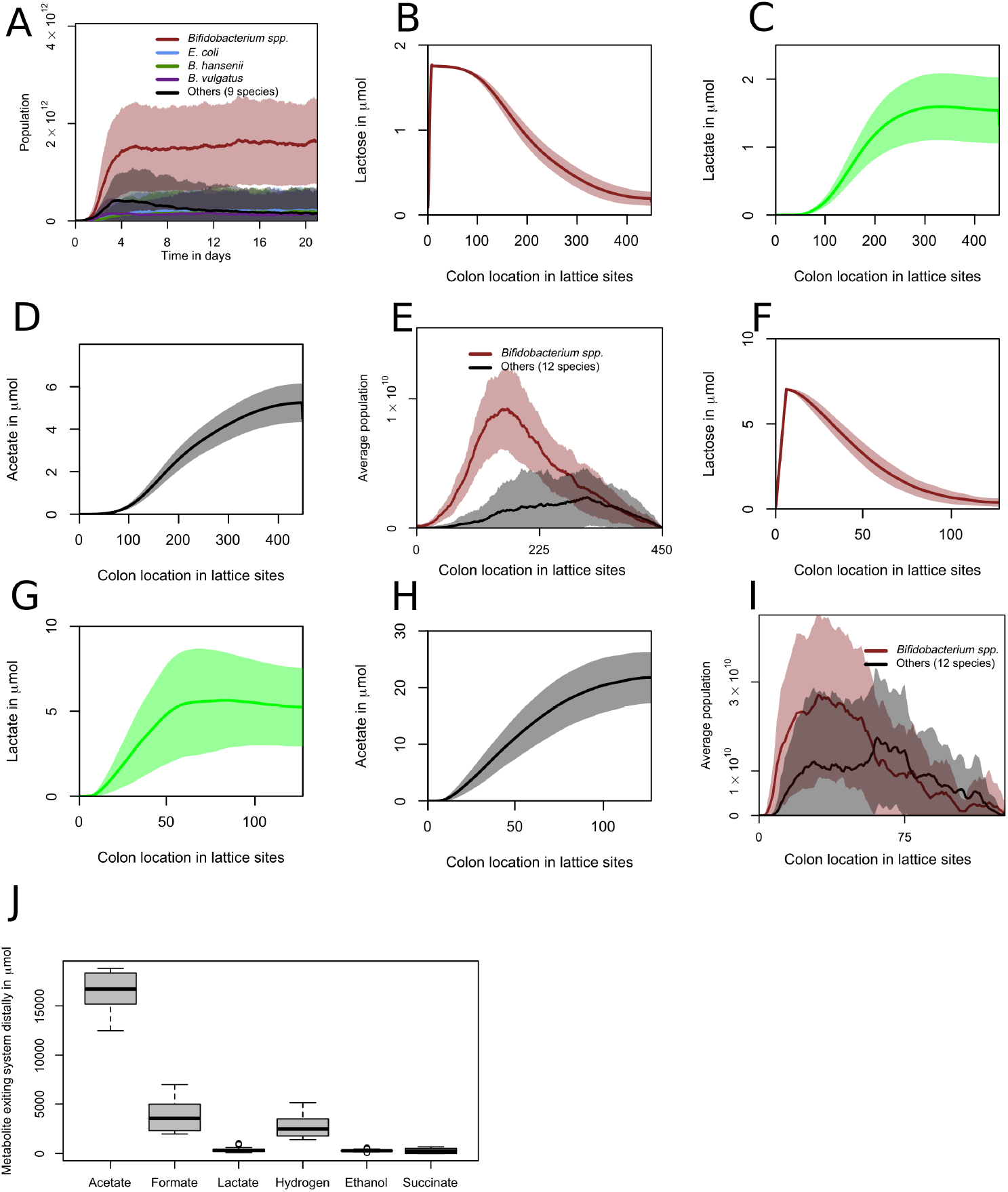
(A) Populations over time with new populations after the initial populations only placed in the first column, with the placement probability (Table 3) increased to 0.01125. Abundances for grouped species over 21 days. One day is 480 timesteps. Curve shows mean value and shaded area one standard deviation over n=30 simulations. (B,C,D) Spatial distribution of lactose, lactate, and acetate, respectively, over the last two days (960 timesteps) of the simulation with a larger lattice. Curve shows mean value and shaded area one standard deviation over n=30 simulations. (E) Abundance per location for grouped species over the last two days (960 timesteps) of the simulations with a larger lattice. Curve shows mean value and shaded area one standard deviation over n=30 simulations. (F,G,H) Spatial distribution of lactose, lactate, and acetate, respectively, over the last two days (960 timesteps) of the simulation with a smaller lattice. Curve shows mean value and shaded area one standard deviation over n=30 simulations. (I) Abundance per location for grouped species over the last two days (960 timesteps) of the simulations with a smaller lattice. Curve shows mean value and shaded area one standard deviation over n=30 simulations. (J) Distribution of metabolites exiting the system distally over the last two days (960 timesteps) of the simulation with diffusion of metabolites increased by a factor of 5. n=30.

**S1 Table. Table of changed or deleted reactions and annotations.csv**

A table of changes made to the AGORA models as a .csv file.

**S2 Table Table of reactions used by *Bifidobacterium longum infantis* in model depending on lactose concentration**

A table containing the flux through each reaction with more than nanomolar flux for *Bifidobacterium longum infantis* populations of 5 · 10^9^ bacteria with access to one lattice site (0.05ml). Water was unlimited, no other metabolites were present. Lactose was present at 0.05, 0.35, or 0.5 µmol per ml.

**S1 Video. Video of a run with no initial oxygen, consisting of a visualisation of the distribution of bacterial species and major metabolites, and a visualisation displaying fluxes between population and metabolite pools. Line width and intensity is proportional to the amount exchanged with the environment over the last 60 timesteps, with a threshold of 0.5 µmol, with no normalization. Metabolite circle size is relative to the most abundant metabolite, with a minimum displayed size of 26 pixels**.

**S2 Video. Visualisations, from left to right, of a run from Fig. 4A, a run from Fig. 4B, and a run from Fig. 4F, displaying population and metabolite pool sizes, and fluxes between populations and metabolites. Line width and intensity is proportional to the amount exchanged with the environment over the last 60 timesteps, with a threshold of 0.5 µmol, with no normalization. Metabolite circle size is relative to the most abundant metabolite, with a minimum displayed size of 26 pixels**.

**S3 Video. Video of a run with an initial 0.1 µmol of oxygen per lattice site, consisting of a visualisation of the distribution of bacterial species and major metabolites, and a visualisation displaying fluxes between population and metabolite pools. Line width and intensity is proportional to the amount exchanged with the environment over the last 60 timesteps, with a threshold of 0.5 µmol, with no normalization. Metabolite circle size is relative to the most abundant metabolite, with a minimum displayed size of 26 pixels**.

**S4 Video. Visualisations, from left to right, of a run from Fig. 5 D, a run from Fig. 5 H, and a run from Fig. 6 A without the enzymatic constraint, displaying population and metabolite pool sizes, and fluxes between populations and metabolites. Line width and intensity is proportional to the amount exchanged with the environment over the last 60 timesteps, with a threshold of 0.5 µmol, with no normalization. Metabolite circle size is relative to the most abundant metabolite, with a minimum displayed size of 26 pixels**.

## CONTRIBUTIONS

P.V.J, M.P., J.M.W.G., and R.M.H.M acquired funding. D.M.V., P.V.J, M.P., J.M.W.G., and R.M.H.M. conceived and planned the simulations. D.M.V. and D.M. wrote software used for the simulations. D.M.V. performed the simulations and analyzed the data. R.S, E.L., J.M.W.G., and R.M.H.M contributed to the interpretation of the results. J.M.W.G., and R.M.H.M. supervised the project. D.M.V. drafted the manuscript. D.M.V., R.S., E.L., J.M.W.G. and R.M.H.M. revised and edited the manuscript.

## ACKNOWLEDGMENTS

This study was financially supported by FrieslandCampina. R.S, E.L., P.V.J., M.P. and J.M.W.G. are currently or were previously employed by FrieslandCampina. The work was carried out in part on the Dutch national e-infrastructure with the support of SURF Cooperative. This work was performed in part using the ALICE compute resources provided by Leiden University.

