## Supplementary figures and images for "A multiscale spatiotemporal model including a switch from aerobic to anaerobic metabolism reproduces succession in the early infant gut microbiota"

### Supplemental Figure 1

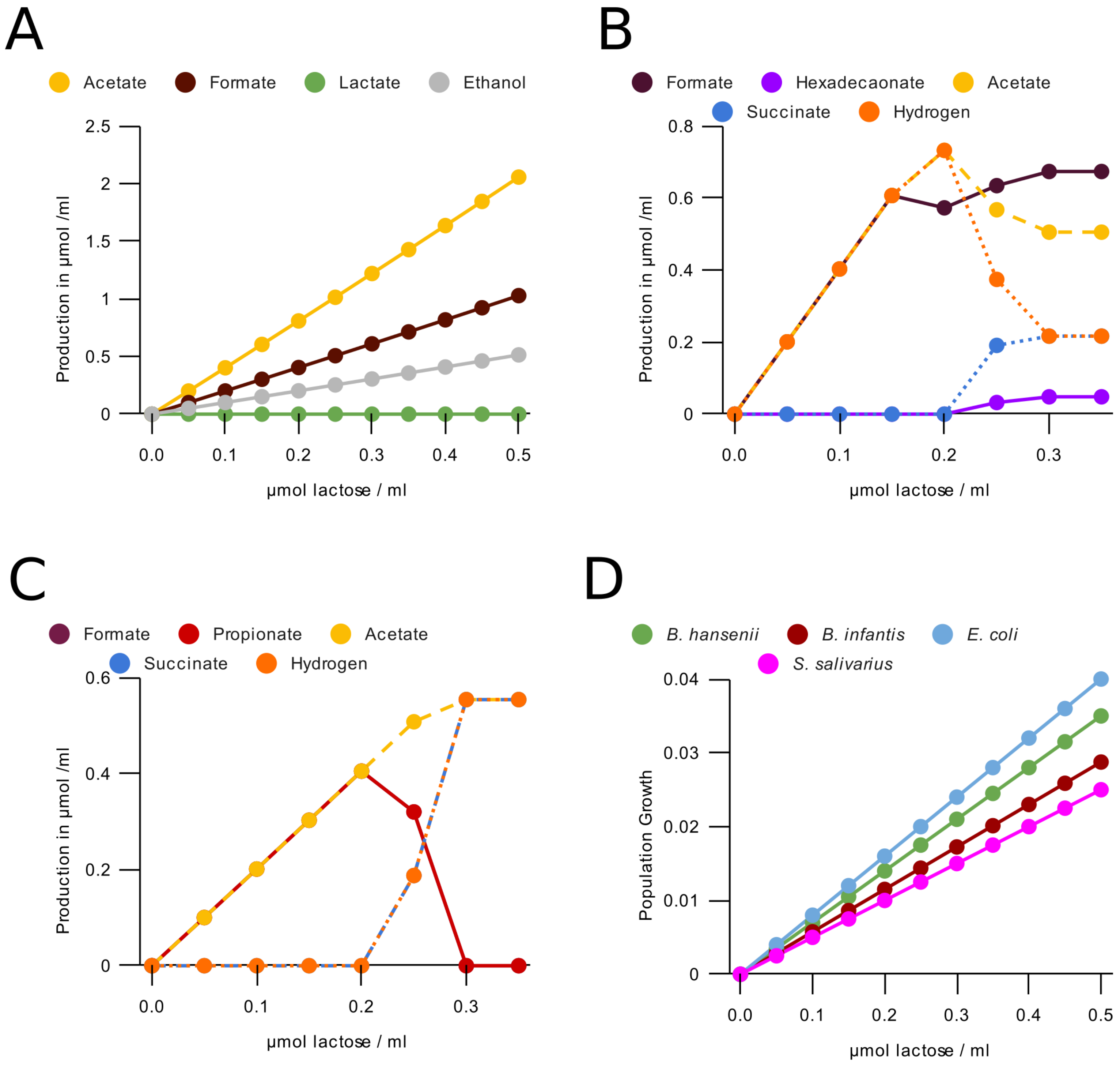

### Supplemental Figure 2

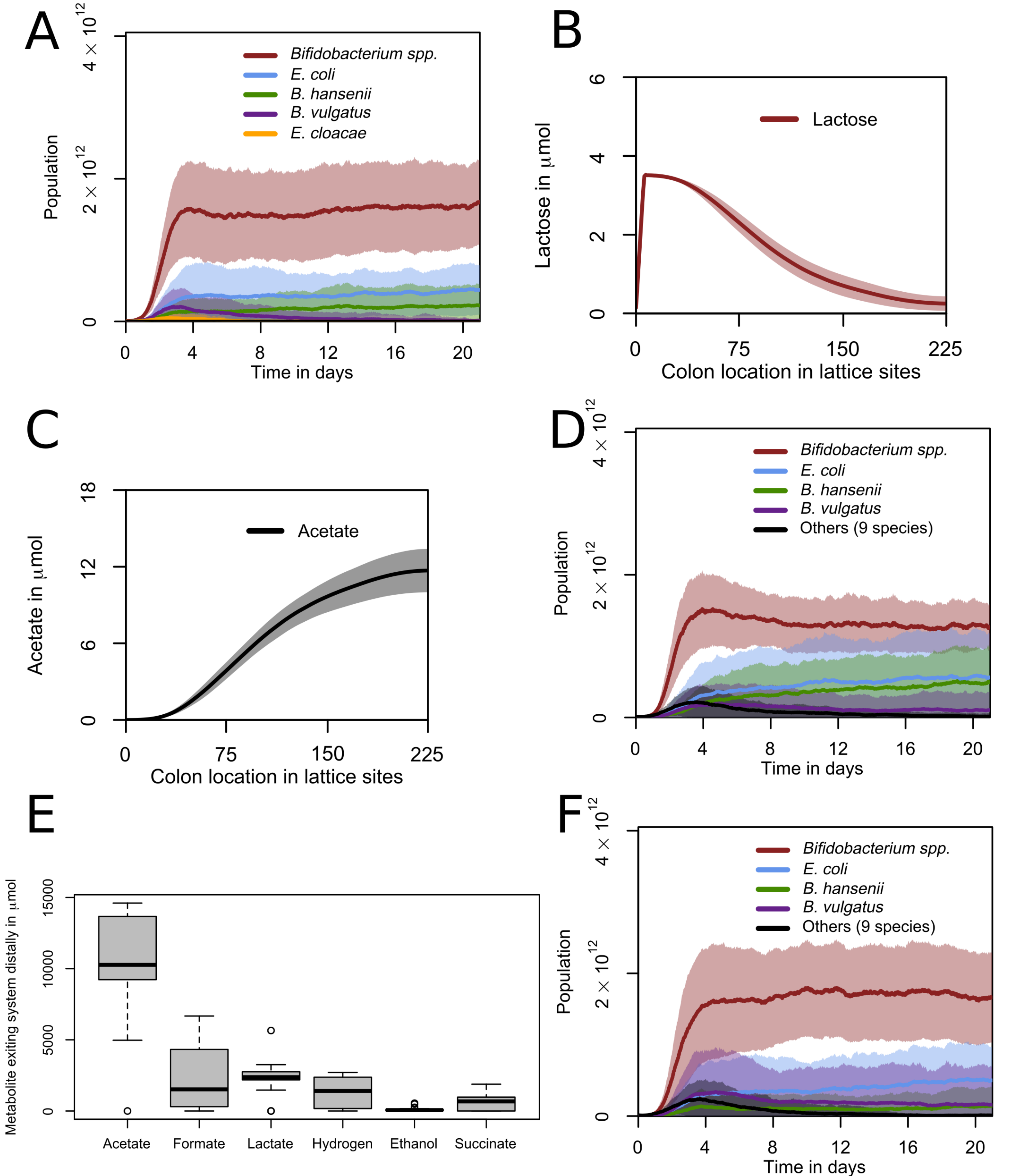

### Supplemental Figure 3

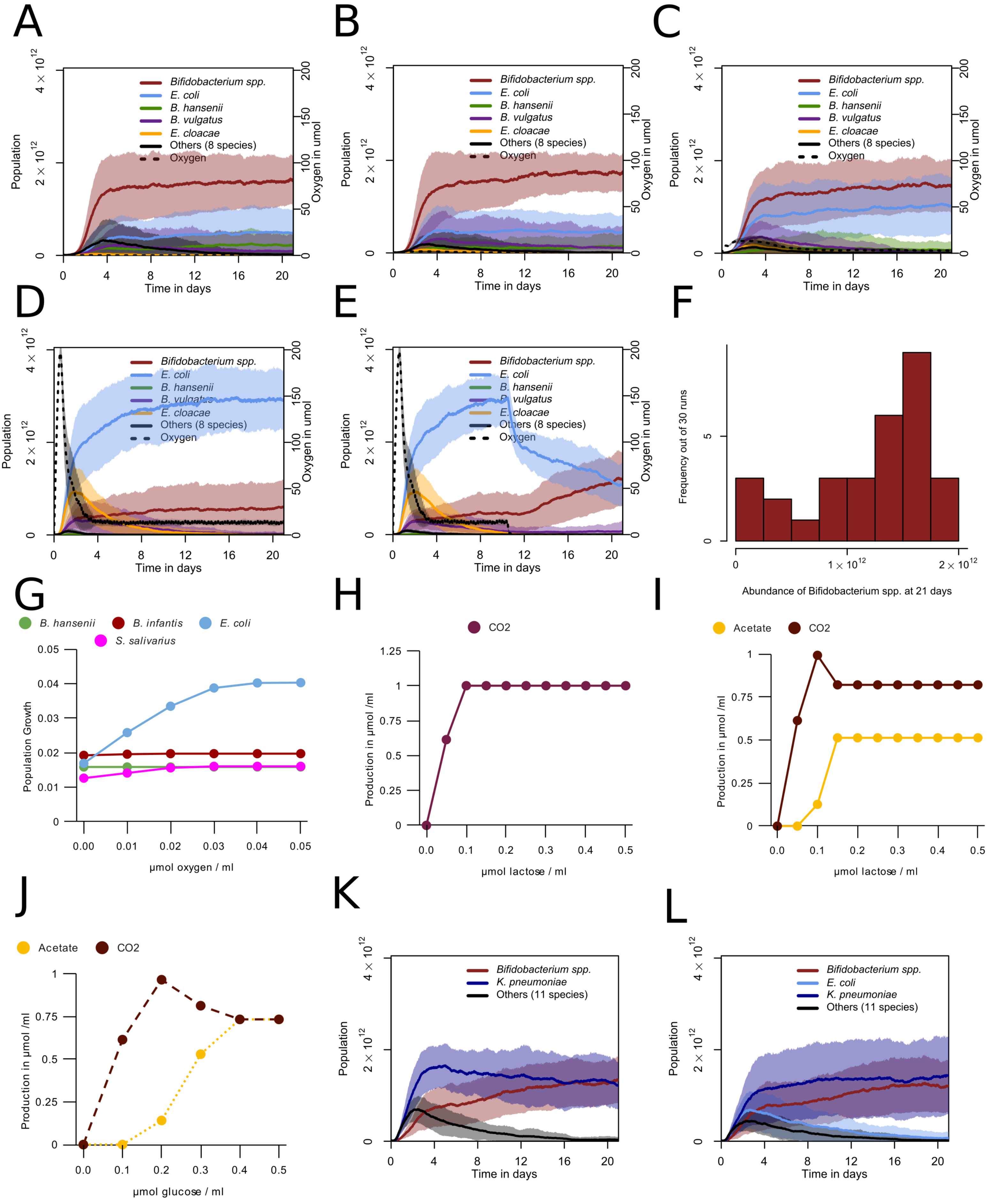

### Supplemental Figure 4

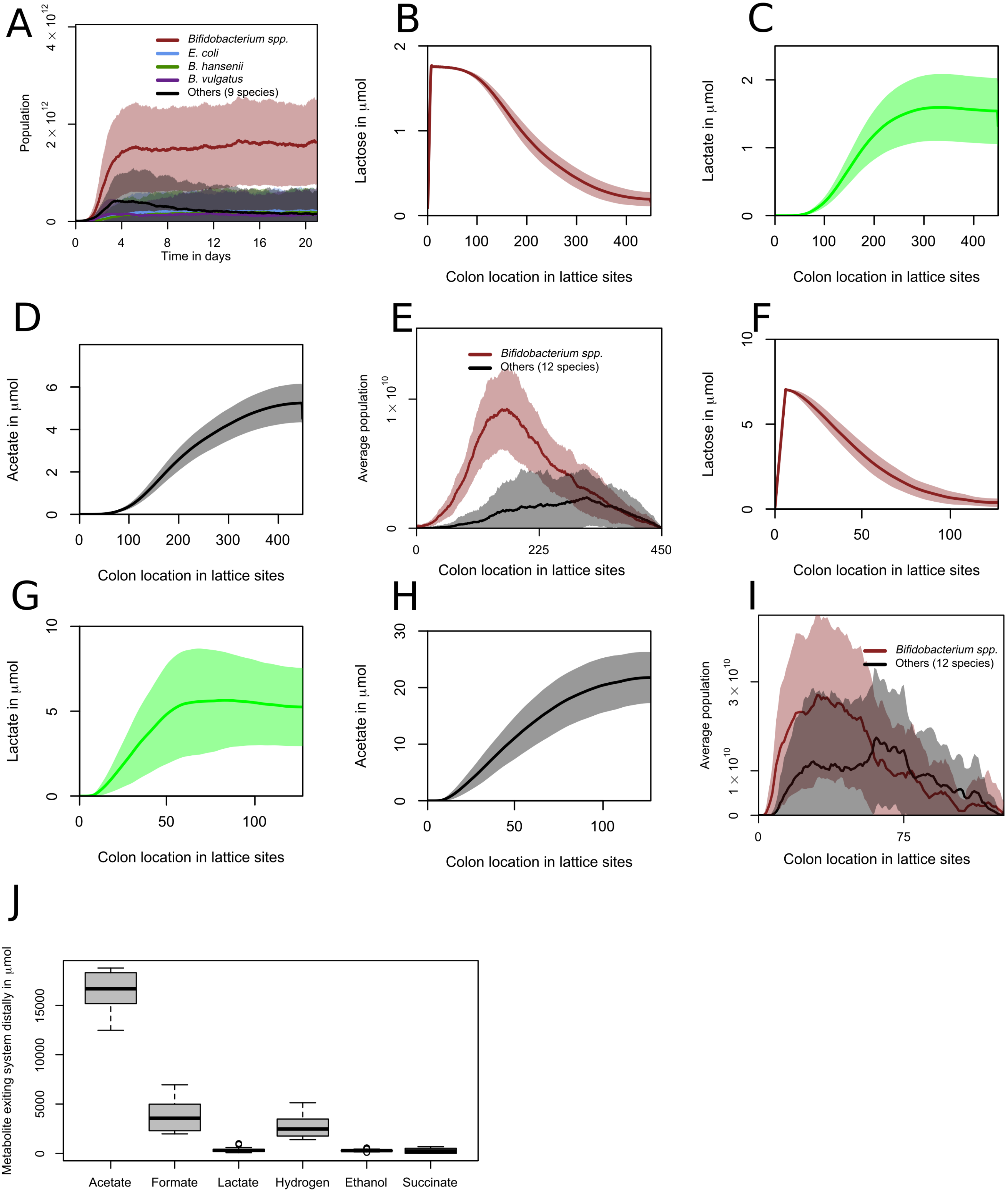
